# Centromeric islands constrain CENP-A domain size and position

**DOI:** 10.64898/2026.10.08.757135

**Authors:** Prachi Patel, Domiziana Trasmondi, Barbara G. Mellone

**Affiliations:** Department of Molecular and Cell Biology, University of Connecticut, Storrs CT, 06269, USA; Institute for Systems Genomics, University of Connecticut, Storrs CT, 06269, USA

## Abstract

Centromere identity is specified epigenetically by nucleosomes containing the histone H3 variant CENP-A, yet centromeres are almost invariably embedded within repetitive DNA, suggesting that underlying DNA sequences contribute to the organization of centromeric chromatin. In *Drosophila melanogaster*, CENP-A primarily occupies retroelement-rich islands flanked by simple satellite DNA, but their functional contribution remains unclear. Here, we delete the ∼50-kb island *Giglio* from centromere 3 while preserving the flanking satellites and residual CENP-A chromatin. Despite removal of the DNA sequence associated with most centromeric CENP-A, centromere identity and overall CENP-A levels are preserved at the native locus, indicating that the island itself is not required for centromere identity. Instead, *Giglio* deletion increases variability in CENP-A chromatin organization, producing highly heterogeneous domain sizes and preferential redistribution of the CENP-A domain into the adjacent *dodeca* satellite. Together, these findings demonstrate that *Giglio* is dispensable for centromere identity but constrains the size and position of the CENP-A domain, establishing centromeric islands as key determinants of CENP-A domain architecture.

## Introduction

Accurate chromosome segregation during mitosis and meiosis depends on centromeres, specialized chromosomal regions that nucleate kinetochores. Kinetochores mediate stable microtubule attachments, ensuring faithful transmission of genetic material to daughter cells (McKinley and Cheeseman 2016). In animals and plants, centromeres are typically embedded within highly repetitive DNA sequences; however, centromere identity is specified not by DNA sequence but by nucleosomes containing the evolutionarily conserved histone H3 variant CENP-A (Palmer et al. 1991), also known as Cid in *Drosophila* (Henikoff et al. 2001) and CenH3 in plants (Talbert et al. 2002). CENP-A nucleosomes are interspersed with canonical H3 nucleosomes (Blower et al. 2002), and their presence is sufficient to establish centromere activity at ectopic loci (Barnhart et al. 2011; Mendiburo et al. 2011; Chen et al. 2014; Palladino et al. 2020). Once assembled, CENP-A chromatin is propagated through a self-templating mechanism supported by dedicated assembly factors, including CAL1 in *Drosophila* and HJURP together with the MIS18 complex in mammals (Dunleavy et al. 2009; Foltz et al. 2009; Chen et al. 2014; McKinley and Cheeseman 2016). If centromere identity and inheritance were fully independent of underlying DNA sequence, this epigenetic framework would predict little constraint on where centromeres are maintained over evolutionary time.

Centromere evolution provides a test for this prediction. Although centromere function is essential and conserved, the DNA sequences underlying centromeres evolve rapidly and diverge even between closely related species. Moreover, centromeres can form *de novo* at ectopic locations, away from ancestral centromeric loci and in the absence of canonical centromere-associated repeats. When such events occur within an individual, they give rise to neocentromeres (Marshall et al. 2008), which can become fixed in populations as evolutionary new centromeres (Ventura et al. 2007). The ability of centromeres to reposition demonstrates that specific DNA sequences are not strictly required for centromere function. Yet despite this epigenetic plasticity, centromere repositioning is rare over evolutionary time, and centromeres are most often stably maintained within large arrays of repetitive DNA across generations within a species. Together, this contrast between flexibility and stability suggests that centromeric DNA, while dispensable for centromere identity, may contribute to centromere positioning long-term.

Experimental studies have shown that centromeric DNA sequences can promote the establishment of CENP-A chromatin under specific conditions. In *S. pombe*, mitotic and meiotic segregation assays using minichromosomes derived from centromeres 1 and 2 demonstrated that the central core and a 2.1 kb fragment of K-type repeat are necessary for centromere function (Baum et al. 1994). Subsequent analyses revealed that the centromere central core DNA promotes elevated H3 nucleosome turnover, favoring CENP-A deposition in fission yeast (Shukla et al. 2018). Human artificial chromosomes (HACs) carrying α-satellite DNA with CENP-B binding motifs can recruit CENP-A and form active centromeres upon introduction into human cells (Ohzeki et al. 2002). These studies demonstrate that specific centromeric DNA sequences can seed centromere formation. However, they primarily address centromere establishment and do not resolve whether underlying DNA sequences are required for centromere maintenance once CENP-A chromatin is formed, or whether they influence the size and stability of pre-existing CENP-A domains.

Beyond contributing to centromere establishment, centromeric DNA sequences have also been implicated in the maintenance and organization of existing CENP-A domains. In *S. pombe*, tRNA genes located at the periphery of centromeres act as boundary elements that prevent heterochromatin from spreading into the central core, thereby protecting centromere identity during meiosis (Scott et al. 2006). In human centromeres, the higher-order repeat (HOR) array that contains CENP-A is consistently the most homogeneous and is flanked by divergent HORs and/or monomeric α-satellite repeats (Altemose et al. 2022; Logsdon et al. 2024). The active array containing CENP-A represents a small portion within the homogeneous HOR and is characterized by a dip in DNA methylation compared to the flanking sequences of the array, known as the Centromere Dip Region (CDR) (Gershman et al. 2022). The CDR delimits the CENP-A region as its expansion through targeted DNA demethylation of flanking alpha-satellite results in the widening of the CENP-A region (Salinas-Luypaert et al. 2025). The CDR has been identified in multiple primates (Makova et al. 2024; Hartley et al. 2025; Yoo et al. 2025). However, how the CENP-A domain is demarcated in organisms lacking a CDR or DNA methylation more generally is unknown.

In a complete human reference genome, the presence of LINE-1 retrotransposons coincides with shifts in DNA methylation, suggesting they may mark the boundary transition (Altemose et al. 2022; Hoyt et al. 2022a). Additionally, human and mouse centromeres contain multiple copies of the 17-bp CENP-B box motif, and CENP-B binding has been implicated in chromosome segregation fidelity in the absence of CENP-A (Dumont and Fachinetti 2017). These observations suggest that centromeric DNA elements can contribute to delimiting and stabilizing centromeric chromatin domains. However, while boundary elements have been functionally tested at endogenous centromeres in fission yeast, the role of CENP-A– associated DNA sequences within metazoan centromeres—particularly in defining the organization, positioning, or stability of established CENP-A domains—remains poorly understood.

*Drosophila melanogaster* provides a uniquely tractable system for dissecting the functional contributions of centromeric DNA within endogenous, established centromeres. Complete assemblies of *D. melanogaster* centromeres revealed that each centromere is organized around a complex DNA “island” enriched in retroelements and flanked by large arrays of simple satellite repeats (Chang et al. 2019). The CENP-A domain spans approximately 101–159 kb, with ∼70% of CENP-A signal localized to the retroelement-rich island and the remaining ∼30% extending into adjacent satellite DNA. An exception is centromere 2, whose island is extremely small (∼2 kb compared to ∼91 kb for other islands), resulting in substantial CENP-A localization within surrounding satellites (Chang et al. 2019).

Although each centromeric island has a distinct sequence organization, none of the DNA elements they contain are exclusive to centromeres. Notably, the non-LTR retroelement *G2/Jockey-3* is the only repeat shared among all five centromere islands (X, 2, 3, 4, and Y) and is highly enriched at centromeres relative to the rest of the genome (Chang et al. 2019; Chabot et al. 2024). The consistent association of CENP-A with retroelement-rich islands—despite the fact that none of the repeats within the islands are unique to centromeres—and the extremely low levels of DNA methylation in this species (Deshmukh et al. 2018)– suggests that the overall organization and composition of these islands may influence CENP-A domain positioning, stability, or boundaries.

Directly testing this possibility requires targeted removal of specific centromeric DNA elements while preserving the surrounding CENP-A domain. Previous whole-centromere DNA deletion studies in human and chicken DT-40 cells resulted in extensive cell death and rare neocentromere formation at ectopic chromosomal locations (Shang et al. 2013; Murillo-Pineda et al. 2021). However, these approaches removed the entire CENP-A domain, precluding analysis of how individual sequence elements influence the organization or behavior of remaining CENP-A chromatin. Moreover, these studies were performed in cultured cells, leaving open whether the same principles apply in a multicellular developmental context, where cellular and tissue-level functions impose additional constraints.

Here, we directly test the functional contribution of centromeric DNA by deleting a major centromeric element—the retroelement-rich island *Giglio* at centromere 3—while preserving the flanking satellites and remaining associated CENP-A chromatin. Using an inducible CRISPR-based chromosome engineering system in developing *Drosophila* tissues, we examined centromere behavior in a multicellular context.

We find that *Giglio* is efficiently deleted in 40–71% of mitotic cells. Despite the removal of the native island, CENP-A reassembles near the deletion site, demonstrating that centromere identity can be re-established in its absence. Notably, analysis of CENP-A-stained metaphase spreads and stretched chromatin fibers reveals that both CENP-A levels and domain size vary substantially in cells carrying the deletion, with *Giglio* removal causing greater variability than CRISPR activity alone. Further analyses using stretched chromatin fibers reveal that, in the absence of *Giglio*, the CENP-A domain shifts into the adjacent *dodeca* satellite and can reach up to 10 times the size of intact centromeres. Together, these findings demonstrate that retroelement-rich islands are not strictly required for centromere identity, but instead constrain both the position and the size of the CENP-A domain, thereby limiting cell-to-cell variability in centromere maintenance.

## Results

### CRISPR-mediated excision of the centromere 3 *Giglio* island and retention of centromere identity

To investigate the role of CENP-A islands in centromere identity and function, we leveraged a *D. melanogaster* centromeric assembly (Chang et al. 2019) along with the established CRISPR system in *Drosophila* ((Gratz et al. 2013); crisprflydesign.org) to design a centromere editing system targeting the centromere 3 island, *Giglio* (**Fig. 1A**). We focused on centromere 3 because its CENP-A domain has been extensively characterized by ChIP-seq and immunofluorescence combined with OligoPaint fluorescence in situ hybridization (IF-FISH) (Chang et al. 2019)In addition, unlike the other centromeres, *Giglio* contains unique sequences flanking both ends of the island that permitted the design of single-guide RNAs (sgRNAs) to excise the entire island through non-homologous end joining (NHEJ)-mediated repair (**Fig. S1**).

**Figure 1.**
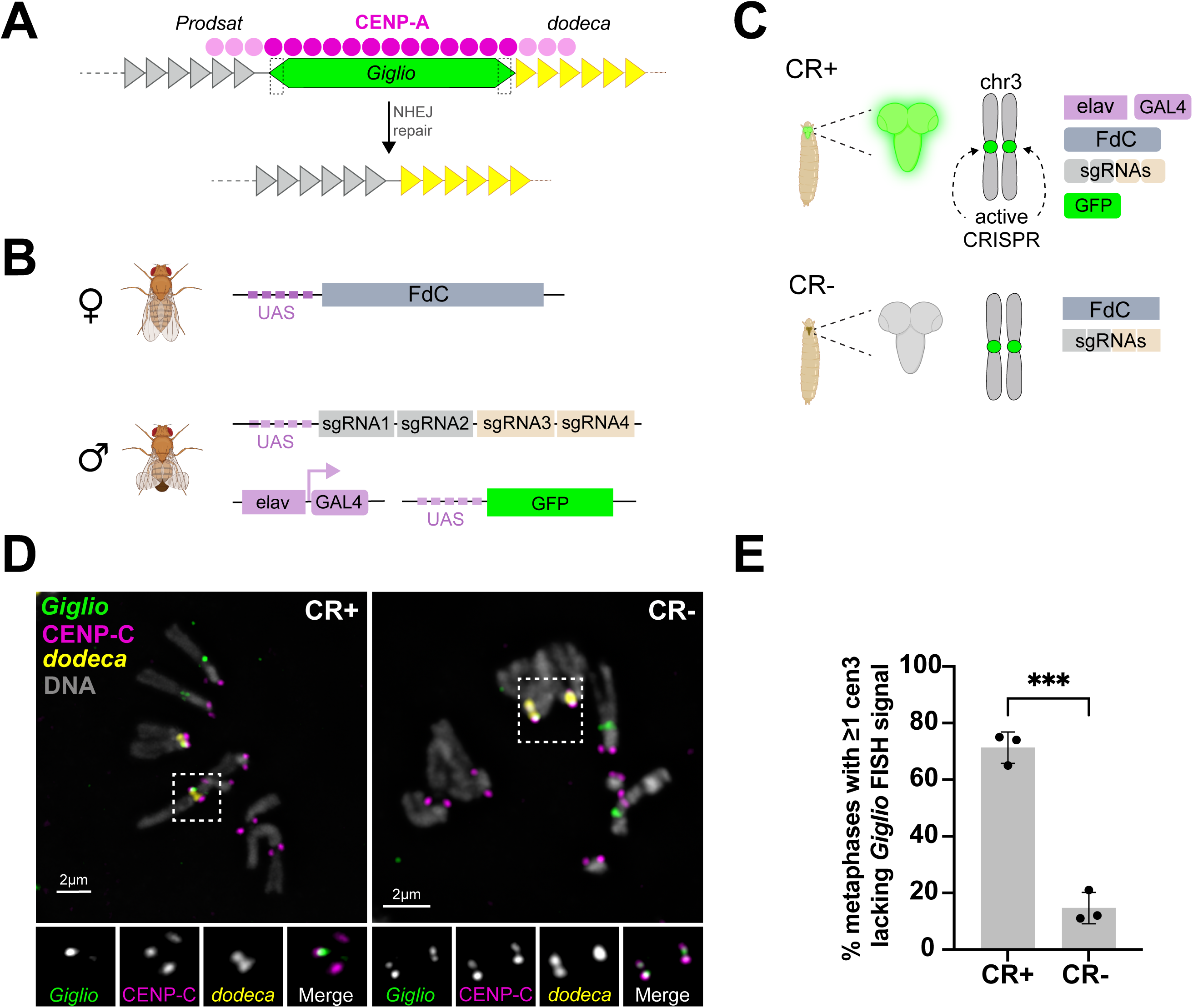
CRISPR-mediated excision of the centromere 3 *Giglio* island and retention of centromere identity. A) Schematic of the centromere 3 (cen3) region (not to scale) showing the CENP-A–enriched *Giglio* island (∼50 kb, green) flanked by *Prodsat* (gray) and *dodeca* (yellow) satellite DNA. Magenta circles indicate CENP-A distribution, with most CENP-A localizing to the *Giglio* island, and a small fraction extending into the flanking satellites (Chang et al., 2019). Dashed boxes mark the unique sequences at each end of the island targeted by the two sgRNA pairs. In the presence of FokI:dCas9 (FdC), paired sgRNAs direct dimerization of FokI nuclease domains, generating DNA double-strand breaks followed by excision of the *Giglio* island through non-homologous end joining (NHEJ). B) Crossing scheme for neuron-specific induction of CRISPR activity. Females carrying UAS-FokI:dCas9 (FdC) on chromosome 2 and an iso-1 chromosome 3 (not shown) were crossed to males carrying a single UAS-4sgRNA transgene encoding sgRNA1-4, together with the pan-neuronal elav-GAL4 driver and a UAS-GFP reporter. sgRNA1-2 (gray) and sgRNA3-4 (tan) correspond to the pairs targeting the 5’ and 3’ ends of the island, respectively. C) The two larval progeny genotypes generated by the cross shown in (B). Squared rectangles indicate transgenes that are present but not expressed; rounded rectangles indicate expressed transgenes. CRISPR-positive (CR+) larvae inherit elav-GAL4, FdC, and the sgRNA transgenes, resulting in neuron-specific CRISPR activity and GFP expression in the brain (green). CRISPR- negative (CR-) siblings inherit FdC and the sgRNAs but lack the GAL4 driver, leaving the system inactive. Both genotypes carry one sgRNA-bearing chromosome 3 and one iso-1 chromosome 3; green dots mark the *Giglio* islands targeted on both homologs. D) Representative mitotic chromosome spreads from third instar larval brains analyzed by immunofluorescence combined with fluorescence in situ hybridization (IF-FISH). Chromosomes were stained with *Giglio* OligoPaint probes (green), a *dodeca* satellite probe (yellow), anti-CENP-C antibodies (magenta), and DAPI (gray). The *dodeca* probe identifies cen3 independently of *Giglio*, allowing chromosome identification following island excision. The CR− metaphase spread contains two *Giglio* signals, whereas the CR+ spread contains one. Insets show magnified views of the boxed centromere 3 region. Scale bars, 2 µm. E) Quantification of *Giglio* deletion efficiency. Bars show the percentage of metaphase spreads containing at least one chromosome 3 lacking *Giglio* FISH. Data represent the mean ± SD from three biological replicates (n = 100 spreads per replicate). **p < 0.01 (unpaired t-test).

To achieve specific DNA cleavage at the target site, we employed the CRISPR/Fok1:dCas9 system (Guilinger et al. 2014), in which two Fok1:dCas9-sgRNA complexes bind adjacent target sequences spaced 13-15 bp apart, allowing dimerization of the fused Fok1 nuclease domains to generate a double-stranded DNA break (**Fig. 1A**). To excise nearly the entire ∼50 kb *Giglio* island, we designed two pairs of sgRNAs, one targeting each end of the island, for a total of four sgRNAs (**Table S2**). The successful excision and NHEJ-mediated of the island is expected to leave behind ∼1.2 kb of island DNA. This residual sequence comprises two *G2/Jockey-3* fragments (100 and 400 bp) at the 5’ end and fragments of the DNA transposon *Transib5* (373 bp) and *G2/Jockey-3* (303 bp) at the 3’ end (**Fig. S1**).

We expressed the sgRNAs and Fok1:dCas9 transgenes using the UAS/GAL4 system to achieve tissue-specific control of expression (**Fig. 1B**). This system produces two types of progeny: CRISPR-positive (CR+), which carry all three transgenes required to activate the CRISPR system, including a UAS-GFP transgene to validate expression in tissues and identify larvae of this genotype, and CRISPR-negative (CR-), which carry the sgRNAs and Fok1:dCas9 but, since they lack the *GAL4* driver, are inactive and thus serve as controls. Both CR+ and CR-genotypes contain one chromosome 3 with the sgRNAs and one from the iso-1 laboratory stock, whose centromeric DNA sequences and CENP-A occupancy were previously identified and characterized (Chang et al. 2019), enabling direct comparison between genotypes (**Fig. 1C** and Methods).

We used the *elav*-GAL4 pan-neuronal driver, which is broadly expressed in larval brain cells and is active in neuroblasts starting from embryogenesis (Berger et al. 2007), to induce expression of our system and analyze it in third instar larval (L3) brain cells, a tissue with high mitotic activity (**Fig. 1C**). To determine the fraction of cells expressing Fok1:dCas9, we quantified UAS-GFP expression in CR+ brains and found that ∼87% of cells are GFP positive whereas CR-brains showed no GFP expression (N=3 brains; n=600 interphase cells; **Fig. S1C-D**).

To assess whether induction of the CRISPR system results in the deletion of *Giglio*, and to determine the consequences for centromere identity, we analyzed mitotic spreads by IF-FISH in larval brains using antibodies against the centromere marker CENP-C (Heeger et al. 2005; Erhardt et al. 2008), *Giglio* OligoPaint probes (Chang et al. 2019) to mark the island, and a probe against *dodeca*, a satellite flanking *Giglio* found only on chromosome 3 (Chang et al. 2019), to aid chromosome identification (**Fig. 1A, C**).

On average, 71% of spreads from CR+ brains contained at least one centromere 3 lacking *Giglio* signal (**Fig. 1D**), indicative of successful island deletion. Importantly, in all instances where centromere 3 lacked *Giglio* FISH signal, CENP-C signal was retained at the deletion site (100%; n=341 *Giglio*-deleted chromosomes; **Fig. 1C**). This result indicates that the centromere was re-established locally, despite deletion of the region that normally contains most CENP-A (Chang et al. 2019). In contrast, approximately 15% of spreads from CR-brains lacked *Giglio* signal on at least one centromere 3. Because these tissues do not express the CRISPR machinery, we interpret this as a technical artifact and use this value as an estimate of the experimental false-positive rate associated with detecting *Giglio* deletion by FISH (i.e., we estimate that 55% of mitotic cells contain true deletions). This frequency was significantly lower than that observed in CR+ brains (71% versus 15%; unpaired t-test, p=0.002). Together, these findings demonstrate successful tissue-specific CRISPR-mediated deletion of the *Giglio* island and retention of centromere identity despite removal of the primary CENP-A-associated DNA region for centromere 3.

### Phenotypes associated with CRISPR activity targeting centromere 3

To establish whether CRISPR-mediated deletion of the *Giglio* island triggers a DNA damage response, we examined the localization of γH2Av, the phosphorylated form of the *Drosophila* homolog of H2AX, which marks sites of double-stranded breaks (Madigan et al. 2002; Mehrotra and McKim 2006). We performed IF on larval brain cells from CR+ and CR-animals using antibodies against CENP-A and γH2Av and analyzed their co-localization in interphase cells (**Fig. 2A**). We found that 13.3% of CR+ L3 brain cells displayed γH2Av signal at the centromere, compared to 3.6% in CR-controls (**Fig. 2B**; t-test, p=0.0093). This ∼3-fold enrichment of γH2Av at centromeres in CR+ brains compared to CR-is consistent with CRISPR-induced double-stranded breaks occurring at the targeted centromeric region. Damage at the centromere in CR-tissues might reflect baseline replication stress, which is known to occur at centromeres (Scelfo et al. 2024). In contrast, the frequency of non-centromeric γH2Av foci was comparable between CR+ and CR-brains (19% vs 18.5%, respectively; t-test, p=0.8396; **Fig. 2C**), suggesting that these foci reflect baseline DNA damage in larval brains under our experimental conditions.

**Figure 2.**
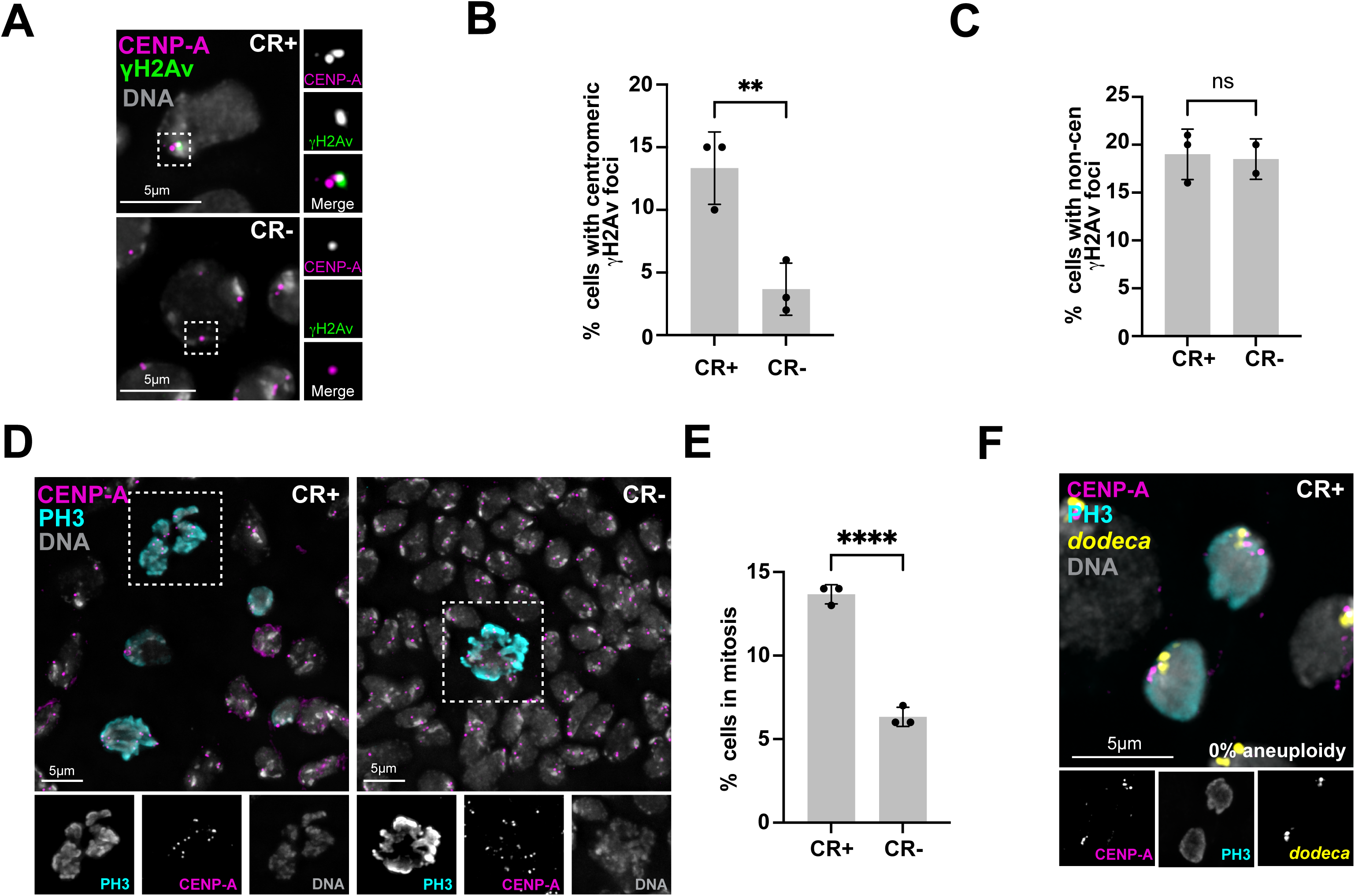
Phenotypes associated with CRISPR activity targeting centromere 3. A) Representative IF images of larval brain monolayers from CR+ (top) and CR- (bottom) animals, stained with antibodies against CENP-A (magenta), γH2Av (green), and DNA (DAPI; gray). Boxed regions show magnified individual channels. γH2Av marks DNA double-stranded breaks. A γH2Av focus overlapping CENP-A is present in the CR+ cell but absent in the CR-cell. Scale bars, 5 µm. B) Percentage of cells with centromeric γH2Av foci (γH2Av overlapping with CENP-A) in CR+ and CR– cells, scored from images as in (A). Each point represents one biological replicate (N = 3 brains; n = 927 cells). Bars show the mean; error bars ± SD. **p < 0.01 (unpaired t test). C) Percentage of cells with non-centromeric γH2Av foci in CR+ versus CR– cells, scored from the same images as in (B). Each point represents one biological replicate (N = 3 brains; n = 927 cells). Bars show the mean; error bars ± SD. ns, not significant (unpaired t-test). D) Representative IF images of L3 larval brain monolayers from CR+ (left) and CR-(right) animals, stained with antibodies against CENP-A (magenta) and phosphorylated histone H3 serine 10 (PH3; cyan); DNA was counterstained with DAPI (gray). PH3 marks mitotic cells. Dash-boxed regions show magnified single-channel images. Scale bars, 5 µm. E) Percentage of PH3-positive (mitotic) cells in CR+ versus CR– cells, scored from images as in (D). Each point represents one biological replicate (N = 3 brains; n = 1043 cells). Bars show the mean, error bars ± SD. ****p < 0.0001 (unpaired t-test). F) Representative IF-FISH image of a CR+ cell undergoing normal anaphase, stained with antibodies against CENP-A (magenta) and PH3 (cyan), a *dodeca* satellite probe (yellow), and DAPI (gray). No lagging chromosomes were observed in any anaphase examined (0%; N = 3 brains; n = 70 anaphases). Insets below show individual channels in grayscale. Scale bars, 5 µm.

To examine whether CRISPR activity affects cell proliferation, we assessed the mitotic index by immunostaining for phospho-serine10 histone H3 (PH3), a mitotic marker (Hendzel et al. 1997; Giet and Glover 2001), in combination with CENP-A on larval brain monolayers (**Fig. 2D**). CR+ brains displayed approximately twice as many PH3-positive cells compared to CR-controls (13.6% in CR+ compared to 6.3% in CR-; t-test, p < 0.0001; **Fig. 2E**) consistent with a mitotic delay, potentially reflecting defective kinetochore/microtubule interactions and activation of the spindle assembly checkpoint (Musacchio and Salmon 2007). However, karyotypic analysis of metaphase spreads revealed no aneuploidy in both CR+ and CR-brains (N = 3 brains, n = 300 spreads; 0% aneuploid in CR+ vs 0% in CR-, p > 0.05). Furthermore, we did not detect lagging chromosomes in anaphase (0%; N=3; n=70; **Fig. 2F**). Although these analyses include all CR+ cells regardless of whether CRISPR-mediated deletion occurred, our IF-FISH analysis indicates that approximately 55% of mitotic cells contain at least one deleted chromosome 3. Thus, if loss of the *Giglio* island severely impaired chromosome segregation, we would expect to detect lagging chromosomes or aneuploid cells at measurable frequencies.

To determine whether the elevated mitotic index correlates with apoptotic cell death, we assessed apoptosis in live whole-mount larval brains using acridine orange (AO) staining, which labels dying cells. CR-brain lobes exhibited baseline AO signal consistent with developmental programmed cell death that occurs during normal larval brain development (Jiang and Reichert 2012). CR+ brain lobes showed elevated AO signal, with AO-positive cells observed as scattered, discrete foci distributed throughout the tissue (**Fig. S2A**). Quantification of the AO signal intensity further confirmed significantly elevated apoptosis compared to CR-(mean ± SD: 15,692 ± 4,688 arbitrary units in CR+ vs 10,191 ± 3,676 arbitrary units in CR-; Welch’s t-test, p = 0.0033; N = 9 CR+ lobes, N = 5 CR-lobes; **Fig. S2B**). Together with the ∼2-fold increase in mitotic index, these findings suggest that CRISPR activity is associated with disrupted cell cycle progression and apoptosis.

To assess whether *Giglio* targeting produces developmental phenotypes, we induced our system using engrailed-GAL4 (broad embryonic expression with segmentation), actin-GAL4 (ubiquitous), and eyeless-GAL4 (developing eye). We scored 100-150 progeny from each cross for eclosion rate or adult eye morphology. None of these drivers produced detectable defects in viability, development, and eye morphology (data not shown). A caveat is that we were unable to assess *Giglio* deletion efficiency in tissues other than larval brains. In larval brains, direct visualization of *Giglio* excision on metaphase spreads indicated deletion in ∼55% of mitotic cells. Deletion efficiency may differ in other tissues could be insufficient to produce detectable phenotypes.

### CRISPR activity and *Giglio* deletion increase variability in CENP-A levels at centromere 3

On centromere 3, approximately 67% of CENP-A localizes to the *Giglio* island, with the remaining CENP-A distributed to the flanking satellites *Prodsat* and *dodeca* (Chang et al. 2019). We therefore predicted that deleting *Giglio* would reduce CENP-A signal intensity, unless more CENP-A is deposited on the flanking satellites to compensate for this loss. To test this prediction, we performed CENP-A IF combined with *Giglio* OligoPaint FISH on mitotic spreads from larval brains, using an acid-free fixation method to retain the presence of CENP-A at centromeres (see Methods; **Fig. 3A**).

**Figure 3.**
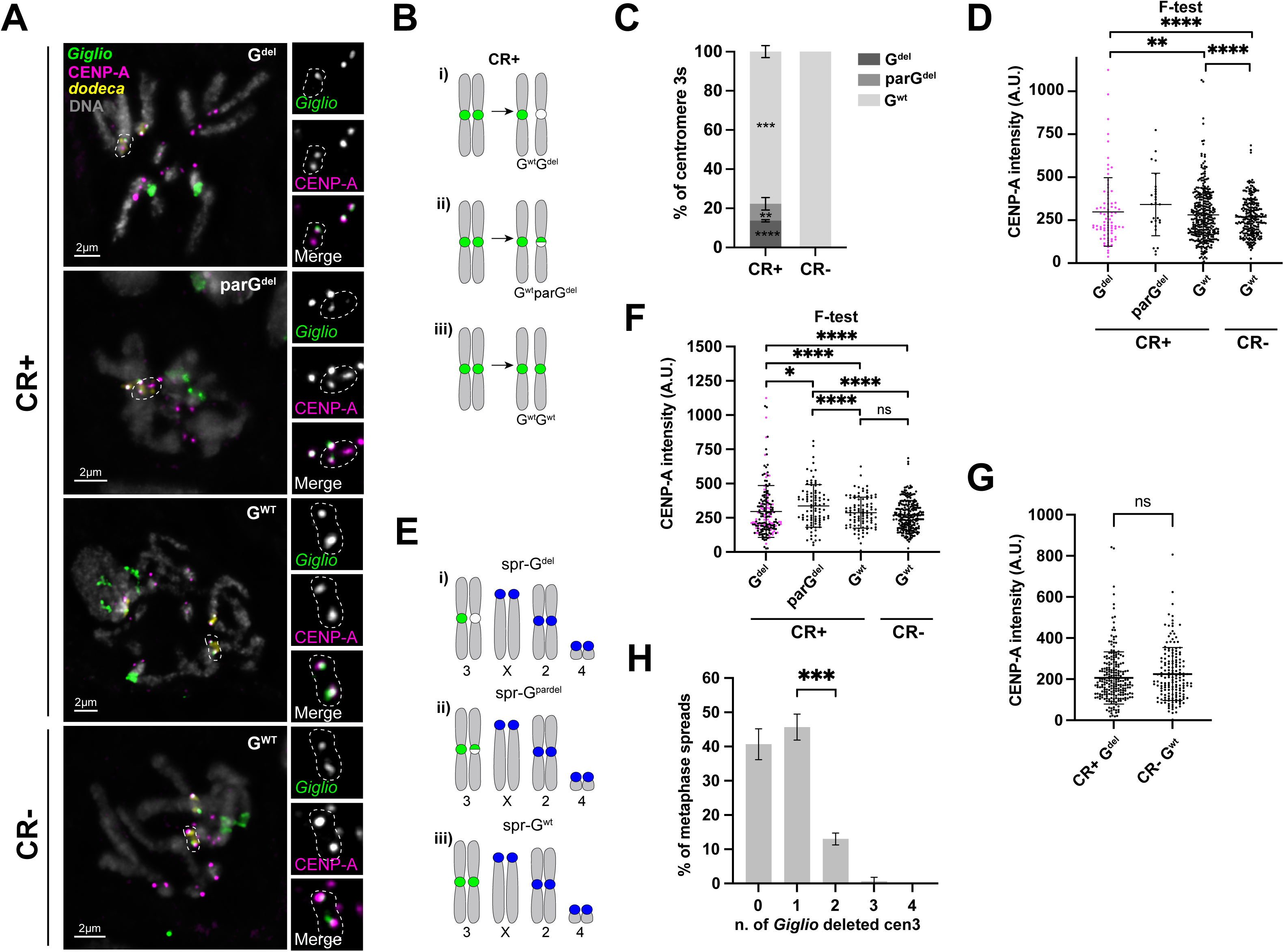
The CENP-A domain size is larger and more variable in response to CRISPR activity and *Giglio* deletion. A) Representative IF-FISH images of metaphase spreads from CR+ (top three) and CR-(bottom) L3 larval brains, illustrating the three centromere 3 classes: G^del^, parG^del^, and G^wt^. Chromosomes were hybridized with *Giglio* OligoPaint probes (green) and *dodeca* satellite probe (yellow), stained with anti-CENP-A antibodies (magenta), and counterstained with DAPI (gray). Insets show higher magnification on the right, with *Giglio* and CENP-A channels in grayscale and the merge in color. Scale bars, 2 µm. B) Schematic of the chromatid-level classification of centromere 3 classes based on the presence, reduction, or absence of *Giglio* FISH signal on each chromatid: (i) G^wt,^ G^del^, one chromatid retaining and one lacking *Giglio*; (ii) G^wt^ parG^del^, one chromatid retaining and one showing reduced *Giglio*; (iii) G^wt^ G^wt^, both chromatids retaining *Giglio*. Green indicates *Giglio* FISH signal, white indicates its absence. C) Percentage of centromere 3’s assigned to each class in CR+ and CR-brains. Bars show the mean of three biological replicates, error bars ± SD (N = 3 brains; n = 300 centromere 3s). Asterisks indicate comparison of each class between CR+ and CR-(unpaired t test): G^del^, ****p < 0.0001; parG^del^, **p < 0.01; G^wt^ ***p < 0.001. D) CENP-A fluorescence intensity (arbitrary units— A.U.) at individual centromere 3’s, grouped by the chromatid-level classification shown in (B): G^del^ (n = 80), parG^del^ (n = 27), and G^wt^ (n = 324) for CR+ and G^wt^ (n = 237) for CR–. Lines show mean ± SD. Statistical comparison shown is F-test for variance: G^del^ vs G^wt^ (**p < 0.01), G^del^ vs CR-(****p < 0.0001), CR+ Gwt vs CR-Gwt (****p < 0.0001). No comparison is shown for parG^del^, for which the p-value could not be determined owing to the small number of data points in this category. E) Schematic of the metaphase spread-level classification of centromere 3: (i) spr-G^del^ are metaphases with one or more G^del^ chromatid, (ii) spr-parG^del^ have one or more parG^del^ chromatid, (iii) spr-G^wt^ have all four centromere 3 chromatids retaining *Giglio*. All centromere 3 in a given cell are quantified and plotted in the corresponding category. Green indicates *Giglio* at centromere 3, blue the centromeres of the remaining chromosomes; one homolog of each chromosome is shown for clarity. F) CENP-A fluorescence intensity (arbitrary units, A.U.) at all centromere 3s, grouped by the metaphase spread-level classification shown in (E): CR+ spr-G^del^ (n = 186), spr-parG^del^ (n = 92), spr-G^wt^ (n = 93), and CR- (n = 237). Lines show the mean ± SD. Statistical comparisons with F-test: spr-G^del^ vs spr-parG^del^, *p < 0.05; spr-G^del^ vs CR+ spr-G^wt^, ****p < 0.0001; spr-G^del^ vs CR- spr-G^wt^, ****p < 0.0001; spr-parG^del^ vs CR+ spr-G^wt^, ****p < 0.0001; spr-parG^del^ vs CR- spr-G^wt^, ****p < 0.0001; CR+ spr-G^wt^ vs CR- spr-G^wt^, ns. G) CENP-A fluorescence intensity (arbitrary units, A.U.) at all non-centromere 3 centromeres (X, 2, 4, and Y) in CR+ spr-G^del^ and CR- spr-G^wt^ metaphase spreads. Lines show the mean ± SD (n = 230 centromeres). Neither the mean nor the variance differed between populations. ns, not significant (unpaired t-test). H) Distribution of *Giglio* deletions among the four centromere 3 chromatids present in CR+ metaphase spreads. Bars show the mean percentage of spreads containing 0, 1, 2, 3, or 4 chromatids lacking *Giglio* FISH signal; error bars ± SD across three biological replicates (N = 3 brains; n = 186 spreads). ***p < 0.001 ([test]).

In contrast to the fixation method used for **Fig. 1**, under these acid-free conditions, all CR-metaphase spreads retained *Giglio* FISH signal, eliminating detectable false-positive *Giglio* loss. Among CR+ samples, we observed three distinct *Giglio* phenotypes with different frequencies: ∼78% of centromere 3 loci showed intact *Giglio* (G^wt^), with FISH signal intensity comparable to that of CR-control; ∼9% were partially deleted (parG^del^), displaying reduced *Giglio* FISH signal compared to CR-samples; and ∼14% appeared fully deleted (G^del^), i.e. lacking detectable *Giglio* FISH signal even after increasing the signal/noise ratio post-imaging (**Fig. 3A-C**). Although only 14% of individual chromatids were G^del^, these deletion events were distributed across ∼44% of metaphase spreads, with an additional ∼25% of spreads containing at least one parG^del^ chromosome, leaving only ∼30% metaphase spreads with intact *Giglio* islands on both chromosome 3 homologs. Importantly, CENP-A signal was retained at all G^del^ centromeres (**Fig. 3A**), i.e., no acentric chromosomes were ever observed (0%; N=3 brains; n=278 spreads). This result indicates that centromere identity is retained, consistent with what we observed in our metaphase spreads stained for CENP-C (**Fig. 1C-D**).

We next quantified the mean CENP-A fluorescence intensity at individual centromere 3s in CR+ and CR-brains. Comparison across centromere 3 categories revealed no significant difference in mean CENP-A intensity across G^wt^, parG^del^, and G^del^ centromeres from CR+ brains relative to G^wt^ centromeres in CR-brains (t-test, p > 0.05; **Fig. 3D**), indicating that total detectable CENP-A levels are maintained following *Giglio* deletion. However, both G^del^ and G^wt^ centromeres from CR+ brains exhibited a higher variance in CENP-A intensity values compared to CR-G^wt^ centromeres (F-test p-value <0.0001; **Fig. 3D**; **Fig. S3**; note that the p-value could not be determined for parG^del^ centromeres due to the low number of data points for this category). The increased variability in G^wt^ centromeres from CR+ brains—which retain intact *Giglio* islands—indicates that CRISPR activity itself, independent of successful deletion, perturbs CENP-A levels.

In our chromatid-level analysis (**Fig. 3D**), the G^wt^ category included centromeres from spreads with confirmed CRISPR activity (i.e., containing at least one G^del^ or parG^del^ chromatid), whereas others came from spreads in which CRISPR-mediated cleavage could not be inferred because all four chromatids appeared intact. In this case, intact chromatids could reflect either successful repair following CRISPR cleavage or the absence of CRISPR activity altogether.

To resolve this ambiguity, we re-categorized centromere 3s according to the status of the entire metaphase spread: spr-G^del^ (spreads containing at least one *Giglio* deletion), spr-parG^del^ (spreads containing at least one partially deleted centromere 3), and spr-G^wt^ (in which all four centromere 3s were intact; **Fig. 3E-F**). This approach accounts for the fact that CRISPR cleavage and repair can occur without producing a detectable deletion.

Using this metaphase spread-level classification, we found that mean CENP-A intensities remained similar across categories, with only spr-parG^del^ showing a modest but significantly higher mean compared to CR-and CR+ spr-G^wt^, (t-test, p = 0.0002 and p = 0.0148, respectively), but not compared to spr-G^del^ (p > 0.05; **Fig. 3F**). In contrast, variance analysis revealed that centromere 3s from spr-G^del^ and spr-parG^del^ metaphases exhibited significantly higher CENP-A intensity variance than those from spr-G^wt^. This difference was observed regardless of whether the centromere 3’s came from CR+ (F-test: spr-G^del^, p < 0.0001; spr-parG^del^, p = 0.0026) or CR-(F-test: both p < 0.0001) animals. The variance between CR+ spr-G^wt^ and CR-spr-G^wt^ was not significantly different (F-test, p > 0.05), indicating that centromeres from spreads with intact *Giglio* behave similarly regardless of genotype. This observation supports the possibility that the CR+ spr-G^wt^ category may be comprised of cells with no ongoing CRISPR activity.

Likewise, variance was modestly but significantly different between spr-G^del^ and spr-parG^del^ spreads (F-test, p = 0.0335), indicating that any level of CRISPR-mediated perturbation generated in a given cell— whether resulting in complete or partial deletion—produces similar effects on CENP-A variability.

To confirm that these effects were specific to centromere 3 targeting rather than a consequence of Fok1:dCas9 or gRNA expression, we quantified CENP-A intensity at all other centromeres (X, 2, 4, and Y combined). Neither mean CENP-A intensity nor variance differed between CR+ and CR-populations at these other centromeres (**Fig. 3G**). Together, these results demonstrate that CRISPR activity with or without successful *Giglio* deletion increases variability in CENP-A levels specifically at centromere 3, resulting in increased CENP-A signal variability.

Lastly, we quantified the number of *Giglio* deletions among the four centromere 3 chromatids present in each mitotic spread. Among CR+ spreads, single-chromosome deletion events were most frequent (∼40%), whereas spreads containing two deletions were less common (13%) and occurred with similar frequency on sister chromatids or on homologous chromosomes. Spreads containing three deletions were rare (0.6%), and no spreads containing four deletions were observed (**Fig. 3H; Fig. S3**).

The predominance of single-deletion spreads, together with the progressive decline in the frequency of higher-order deletion events and the absence of spreads containing four deletions, is consistent with CRISPR activity occurring throughout embryonic and larval development, resulting in the continual generation of new detectable *Giglio* deletion events within the mitotic population. Because we also observed increased apoptosis in CR+ brains, cell loss may contribute to the underrepresentation of spreads containing multiple *Giglio* deletions. Notably, deletions affecting both sister chromatids are consistent with *Giglio* being removed prior to DNA replication, whereas deletions affecting homologous chromosomes suggest independent events within the same cell. Together, these data indicate that CRISPR-mediated *Giglio* deletions are frequent but are predominantly observed as single-chromatid events in mitotic cells.

### The CENP-A domain size is larger and more variable in response to CRISPR activity and *Giglio* deletion

Our data so far shows that, upon *Giglio* deletion, centromere 3 is re-established at or near the deletion site, and that CENP-A intensity is more variable in spreads where CRISPR is active, as inferred from the presence of full or partial deletion of *Giglio* from one or more centromere 3 (**Fig.3**). The CENP-A domain of centromere 3 measures approximately 10 µm on stretched chromatin fibers, corresponding to ∼101 kb, with 67% of CENP-A signal overlapping with the *Giglio* island (Chang et al. 2019). Since *Giglio* and the retroelement-rich islands in general appear to be preferred sites for CENP-A assembly, we hypothesized that its removal might destabilize CENP-A occupancy and result in perturbation of the domain’s length.

To investigate changes in CENP-A domain size following *Giglio* deletion, we generated stretched chromatin fibers from CR+ and CR-larval brains and employed IF-FISH to simultaneously visualize CENP-A, the *Giglio* island, and the *dodeca* satellite (**Fig. 4A**). The *dodeca* satellite serves as a marker for identification of centromere 3 in the absence of *Giglio*, as it is located exclusively adjacent to the *Giglio* island on chromosome 3 (**Fig. 1A**), unlike *Prodsat*—the satellite flanking the opposite side of *Giglio*— which is also located at the pericentromere of chromosome 2 (Chang et al. 2019). We analyzed long, single chromatin fibers displaying CENP-A colocalization with *Giglio* and/or *dodeca*. In CR-control samples and G^wt^ fibers from CR+ brains, CENP-A predominantly colocalized with the *Giglio* island, with additional staining in the flanking *dodeca* region (**Fig. 4A**), consistent with previous reports (Chang et al., 2019). We also note that CENP-A is visible along a short stretch in the opposite *Giglio*-flanking region, where *Prodsat* is located, in a subset of fibers, consistent with moderate CENP-A enrichment in this region (3/30 fibers for CR-and 2/30 for CR+ G^wt^; **Fig. 4A**).

**Figure 4.**
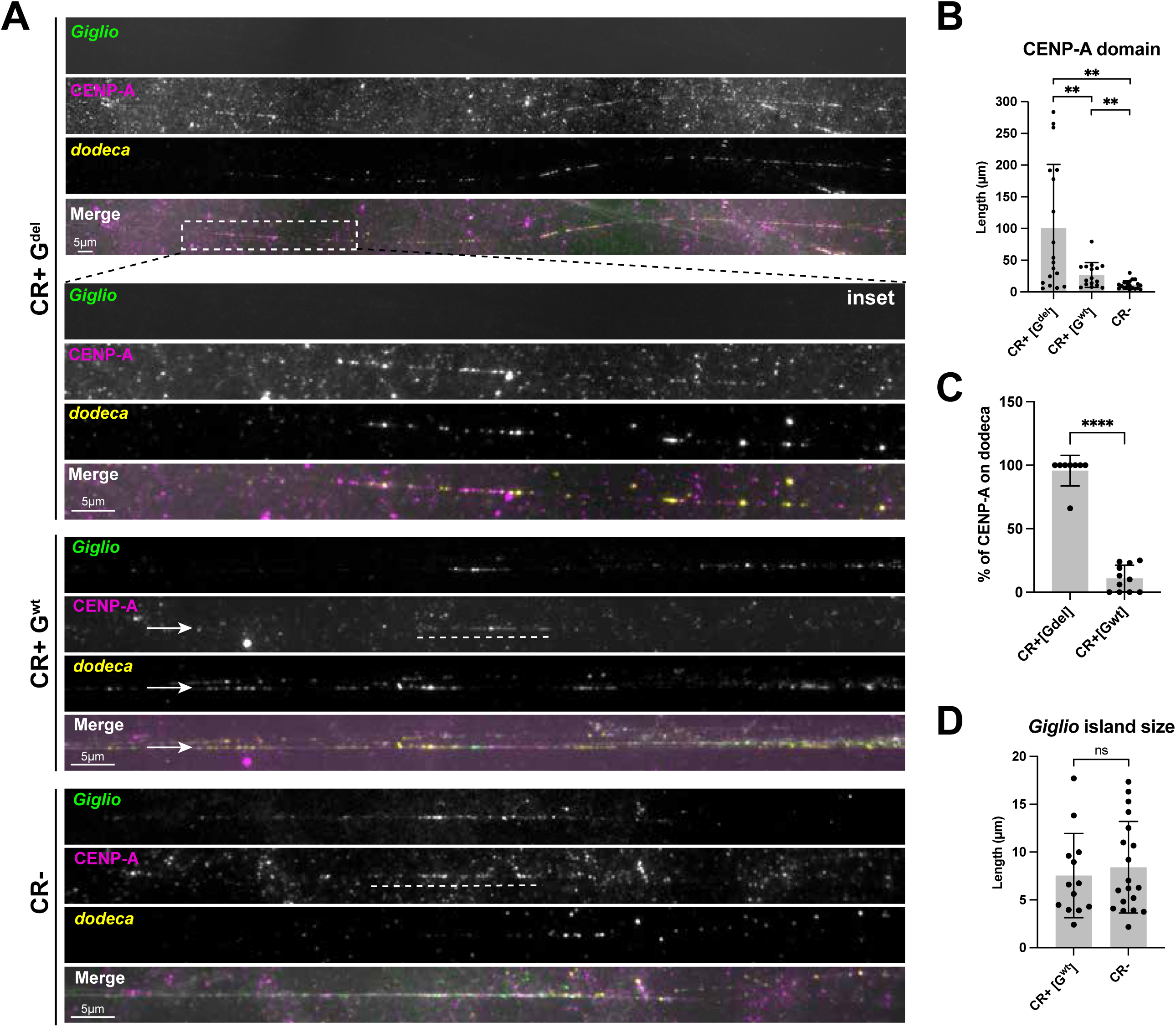
CENP-A domain size increases in size and variability following Giglio deletion. A) Representative IF-FISH images of extended chromatin fibers from CR+ brains (G^del^ and G^wt^) and CR– brains. *Giglio* OligoPaint FISH signal is shown in green, CENP-A IF in magenta, *dodeca* in yellow, and DNA (DAPI) in gray. The dashed box on the CR+ G^del^ fiber indicates the region shown enlarged in the inset below. The white dashed line indicates the CENP-A domain. Scale bars, 5 µm. B) Quantification of CENP-A domain length (μm) in extended chromatin fibers: CR+ G^del^ (n = 20 fibers), CR+ G^wt^ (n = 18 fibers), and CR- (n = 20 fibers). Bars show the mean, errors bars ± SD. **p < 0.01 (unpaired t-test). C) Percentage of the CENP-A domain overlapping the *dodeca* satellite on individual fibers from CR+ brains: G^del^ (n = 8 fibers) and G^wt^ (n = 11 fibers). Bars show the mean, error bars ± SD. ****p < 0.0001 (unpaired t-test). D) Length (µm) of the *Giglio* island FISH signal measured on extended chromatin fibers from CR+ G^wt^ (n = 13 fibers) and CR- brains (n = 19 fibers). Bars show the mean, error bars ± SD. Comparable island lengths indicate that fiber stretching was consistent between populations. ns, not significant (unpaired t-test).

To assess the impact of *Giglio* deletion, we categorized centromere 3 fibers from the CR+ population based on the presence (G^wt^) or absence (G^del^) of the island as assessed by FISH. We did not include a parG^del^ category because it was difficult to distinguish whether a fiber with a short domain containing *Giglio* signal represented a partially deleted island or non-specific probe hybridization. There is also a short region with *Giglio* OligoPaints FISH signal found within the flanking *dodeca* block of chromosome 3 (Chang et al. 2019) that complicates the interpretation of such fibers.

In G^del^ fibers, the CENP-A domain displayed substantial size heterogeneity, ranging from 5.5 µm to 284 µm in length. In contrast, fibers with intact *Giglio* from CR-brains displayed CENP-A domains between 3.7 µm and 30 µm long, in line with previous measurements (**Fig. 4A-B and Fig. S4**; (Chang et al. 2019). The mean CENP-A domain length in G^del^ fibers (100.5 µm) was significantly higher than that of G^wt^ fibers from both the CR+ (26.9 µm; t test p=0.0124) and the CR-populations (11.3 µm; t test p=0.0015; **Fig. 4B**). Consistent with the broader size distribution, the variance in CENP-A domain length was significantly greater in the G^del^ population than in G^wt^ centromeres from both CR+ (F-test, p = 0.0001) and CR-populations (F-test, p < 0.0001). These results show that loss of *Giglio* is specifically associated with increased variability in CENP-A domain length, suggesting perturbed control of CENP-A localization.

We also observed that the CENP-A domain predominantly associated with *dodeca*. To quantify this apparent distribution, we measured the proportion of each CENP-A domain that overlapped *dodeca* on individual fibers from CR+ brains (**Fig. 4C**). In G^del^ fibers, essentially the entire CENP-A domain resided within dodeca (95.75%; n = 8 fibers), with complete overlap observed in all but one fiber. In contrast, G^wt^ fibers from CR+ brains retained the wild-type arrangement with only a minor fraction of the domain extending into *dodeca* (10.9%; n = 11 fibers; t test, p<0.0001). Because *Giglio* normally accommodates ∼67% of CENP-A (Chang et al. 2019), this near-complete association with *dodeca* indicates preferential shift of the CENP-A domain. Together with the increase in domain length (**Fig. 4B**), these results indicate that *Giglio* deletion both expands the CENP-A domain and shows a preference for *dodeca* satellite.

Although all G^wt^ fibers retain an intact *Giglio* island, the mean CENP-A domain length and its variance were significantly greater in CR+ than in CR− brains (26.9 µm vs. 11.3 µm; **Fig. 4B**; t test p=0.0072; F-test, p < 0.0001). Because chromatin fibers are isolated from a mixed population of cells, we cannot determine which G^wt^ fibers originated from cells in which CRISPR activity had occurred. Nevertheless, this observation is consistent with our mitotic chromosome analysis (**Fig. 3E–F**), which showed increased CENP-A intensity variability at intact centromere 3s from spreads containing *Giglio* deletion events, and indicates that increased CENP-A intensity variability reflects increased variability in CENP-A domain size.

To rule out the possibility that differences in CENP-A domain size between G^wt^ fibers from CR+ and CR-brains result from differential stretching of the chromatin fiber samples, we compared the length of the *Giglio* island itself between these populations, with the expectation that island size should be consistent across both CR- and CR+ genotypes in non-deleted chromosomes. Indeed, the *Giglio* island FISH signal exhibited similar mean lengths and size distributions in both populations (**Fig. 4D**), demonstrating that chromatin fiber stretching was consistent across samples. This analysis supports the conclusion that increased size heterogeneity of the CENP-A domain is a consequence of CRISPR activity and *Giglio* deletion at centromere 3.

Our comparisons distinguish the effects of CRISPR activity from those of *Giglio* deletion on the CENP-A chromatin domain. In the absence of detectable *Giglio* deletion, CENP-A domains in CR+ fibers were moderately larger than those in CR-controls. Upon *Giglio* deletion, however, CENP-A domains expanded dramatically and exhibited substantially greater variability (**Fig. 4B**). Together with our measurements of CENP-A fluorescence intensity (**Fig. 3**), these findings indicate that maintenance of total CENP-A levels following *Giglio* deletion is accompanied by substantial expansion and increased heterogeneity of the CENP-A chromatin domain.

### Analysis of CENP-A occupancy across *Giglio* and the surrounding satellites

Chromatin fibers provide single-chromosome resolution and allow G^del^ and G^wt^ centromeres to be distinguished directly, but they sample a limited number of single chromosomes. To determine whether the changes in CENP-A domain size and position observed on fibers are also detectable as changes in read coverage, we profiled CENP-A genomically using CUT&Tag (Kaya-Okur et al. 2019). An important consideration for this approach is that *Giglio* deletion is mosaic: only ∼14% of centromere 3 chromatids lack detectable *Giglio* by FISH in CR+ larval brain cells (**Fig. 3C**), so any population-based assay performed on whole larval brains averages CENP-A signal across a majority of intact centromere 3s.

We performed CENP-A CUT&Tag on brains from CR+ and CR-larvae, using IgG as a negative control. Both genotypes carry identical heterozygous chromosome 3s (UAS-sgRNAs/+Iso1; **Fig. 1B**), so differences between them are attributable to CRISPR activity and *Giglio* deletion. Reads were mapped to the heterochromatin-enriched *D. melanogaster* genome assembly (Chang et al. 2019), normalized to counts per million (CPM), and visualized as log2(CENP-A/IgG) coverage across the annotated centromere 3 contig, together with MACS2 peak calls (**Fig. 5A**).

**Figure 5.**
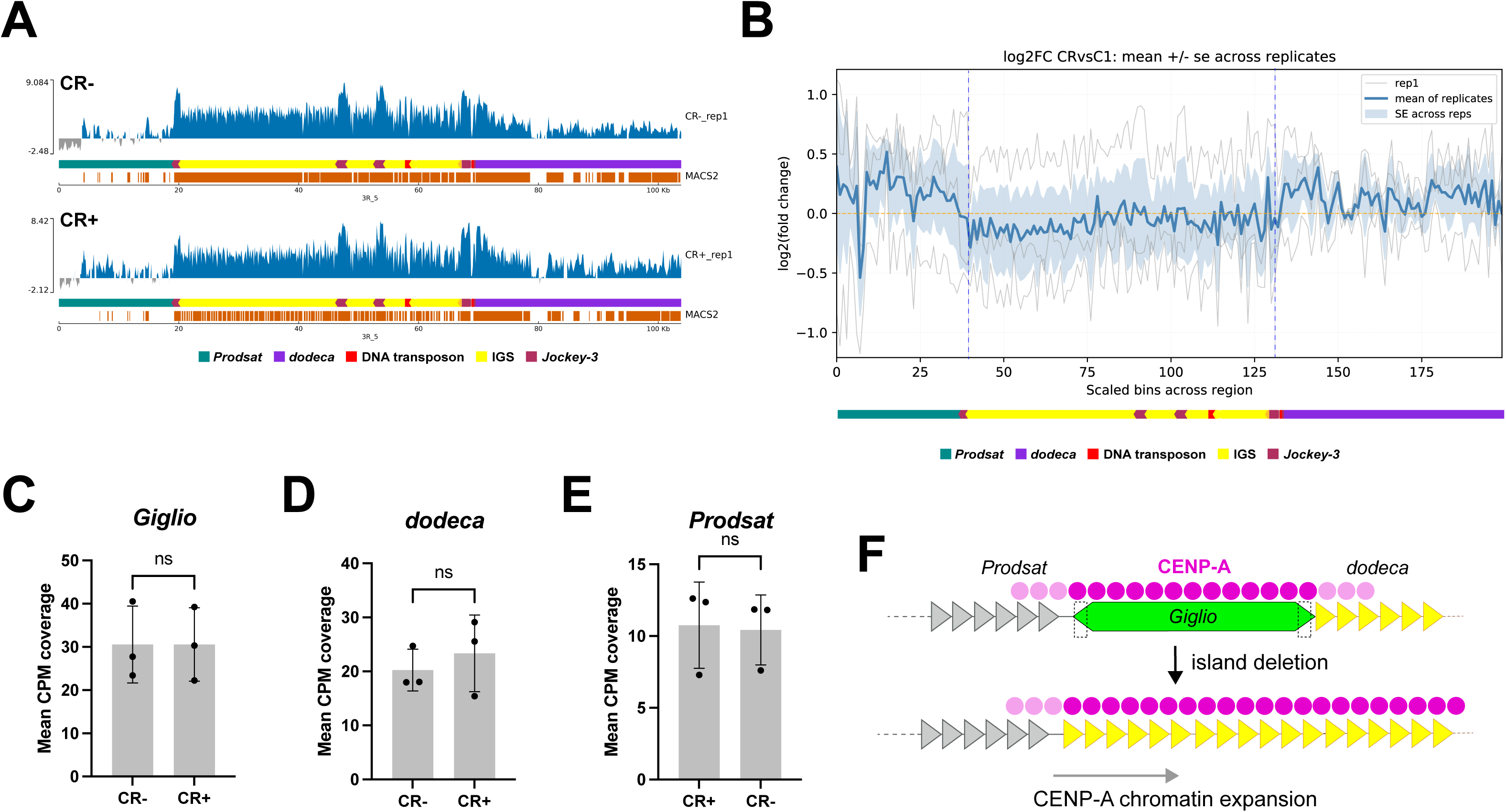
Analysis of CENP-A occupancy across Giglio and the surrounding satellites. A) CENP-A CUT&Tag coverage across the centromere 3 contig in CR- (top) and CR+ (bottom) larval brains, shown as log2(CENP-A/IgG) read coverage for one representative replicate per population. Positive values are shown in blue and negative values in gray. The annotation track indicates *Prodsat* (teal), the *Giglio* island (yellow, with retroelement annotations in magenta and red), and *dodeca* (purple). MACS2 peak calls for each sample are shown below the annotation (orange). B) Log2 fold change in CENP-A coverage between CR+ and CR- across scaled bins spanning *Prodsat*, *Giglio*, and *dodeca*. Thin gray lines show individual replicates, the blue line the mean across replicates, and shading ± SE across replicates. The horizontal dashed line marks a log2 fold change of zero; vertical dashed lines indicate the sgRNA cut sites at the island boundaries. C-E) Mean CPM coverage of CENP-A CUT&Tag reads across *Giglio* (C), *dodeca* (D), and *Prodsat* (E) in CR- versus CR+ larval brains. Each point represents one biological replicate (n = 3). Bars show the mean, error bars ± SD. ns = not significant (unpaired t-test). F) Model for the reorganization of the CENP-A domain upon deletion of the centromeric island via CRISPR. Upon loss of the island and its associated CENP-A chromatin, new CENP-A is deposited to the flanking satellite, with a preference for *dodeca*. The size of the CENP-A domain changes from being consistent across cells to being misregulated, resulting in CENP-A domains of highly variable size, including significantly expanded ones.

To directly compare CENP-A distribution between genotypes, we generated log2 fold-change profiles across the centromere 3 region, in which positive values indicate relative enrichment in CR+ samples and negative values indicate relative depletion compared to CR-(**Fig. 5B**). We observed a consistent directional pattern across the region, with most peaks across *Prodsat* and *dodeca* shifting above zero, whereas most peaks across the *Giglio* island shifted below zero. Notably, the transitions between relative enrichment and relative depletion did not fall at arbitrary positions within the region but coincided with the boundaries of the island, where the sgRNA target sites used to excise *Giglio* are located (**Fig. 5B** and **Fig. S1A**). These profiles reveal relative depletion of CENP-A reads across the island concurrent with relative enrichment across both flanking satellites in CR+ compared to CR-brains, delimited by the engineered break sites. This coincidence is consistent with the redistribution being a direct consequence of island excision rather than a general difference between the two populations. Because *Giglio* deletion is mosaic, the magnitude of these shifts is expected to be small relative to the variability between replicates.

To ask whether this redistribution was also detectable as a change in absolute CENP-A occupancy, we quantified mean CPM coverage separately across *Giglio*, *dodeca*, and *Prodsat* in each replicate (**Fig. 5C-E**). Coverage across *Giglio* was indistinguishable between genotypes (mean ± SD; n = 3 biological replicates; ns, unpaired t test; **Fig. 5C**). Coverage across *dodeca* was modestly higher in CR+ brains, in the direction predicted by the differential profiles and by the fiber measurements, but this difference was not significant and was accompanied by greater replicate-to-replicate spread (**Fig. 5D**). Coverage across *Prodsat* was similarly unchanged (ns; **Fig. 5E**). These measurements did not resolve a significant difference in CENP-A occupancy in any region, though the largest change was an increase across *dodeca* in CR+ brains.

To ask whether CENP-A occupancy was altered at individual enriched sites rather than across entire regions, we compared the intensities of individual CENP-A peaks called by MACS2 within *Giglio*, *dodeca*, and *Prodsat* (**Fig. S5A-C**). Peak intensities were broadly distributed in both populations, with a small number of high-intensity peaks in each region, and did not differ significantly between CR+ and CR- in any of the three regions (ns, unpaired t test). This comparison is restricted to sequences at which CENP-A enrichment was sufficient for a peak to be called, so it reports the strength of retained peaks rather than changes in the distribution. Thus, neither regional coverage nor the intensity of individual CENP-A peaks was detectably altered by CRISPR activity or deletion at the population level.

To determine what fraction of the population carried the *Giglio* deletion in these mosaic brains, we performed whole-genome shotgun sequencing of CR+ and CR-larval brains and compared read depth across centromere 3 (**Fig. S5D**). Coverage across the interval between the two sgRNA cut sites was comparable between populations, and the log2 coverage ratio remained close to zero across the island as well as across the flanking satellites. Because excision removes *Giglio* entirely from the locus, a homogeneous deletion would produce a marked reduction in read depth across it. Its absence is consistent with the mosaicism observed cytologically (**Fig. 3C**).

Together, these analyses are consistent with a redistribution of CENP-A away from the island and into the flanking satellite sequences following *Giglio* deletion, but do not resolve the magnitude of this change.

This is expected given the mosaic nature of our system: because only a minority of centromere 3 copies in the brain carry a full deletion (**Fig. 3C**), CUT&Tag signal is dominated by intact centromere 3s contributing unchanged coverage at *Giglio*, *dodeca*, and *Prodsat*, and reads cannot be attributed to deleted versus intact chromosomes. By contrast, chromatin fibers allow deleted centromeres to be identified and measured individually, and indicate that the redistribution occurs preferentially on *dodeca* in the absence of the island (**Fig. 4C**; **Fig. 5F**). We therefore interpret the fiber measurements as reporting the behavior of *Giglio* deleted centromere 3s, and the CUT&Tag profiles as a diluted, population-averaged view of the same redistribution.

## Discussion

In this study, we used an inducible CRISPR/Fok1 system to investigate the effects of deleting the ∼50 kb retroelement-rich *Giglio* island from centromere 3 in *D. melanogaster* while preserving the flanking satellite DNA and residual CENP-A chromatin. Despite removal of the DNA sequence associated with the majority of CENP-A on centromere 3, its identity was maintained at the native locus, indicating that *Giglio* is not required for centromere maintenance. At the same time, *Giglio* deletion altered CENP-A domain organization, increasing cell-to-cell variability in CENP-A levels, expanding domain size, and shifting CENP-A preferentially into the adjacent *dodeca* satellite. Together, these findings indicate that centromeric retroelement-rich DNA is dispensable for maintaining centromere identity but plays an important role in constraining the position and size of the CENP-A domain.

Despite mosaicism in CR+ brains, targeted deletion of *Giglio* enables direct testing of the contribution of centromeric DNA to centromere identity and function *in vivo.* Following *Giglio* deletion, centromere 3 could in principle be lost, repositioned, or re-established locally from residual CENP-A chromatin. Our results strongly support the latter outcome.

Retention of CENP-A and CENP-C at all deleted centromeres indicates that centromere identity is maintained locally despite removal of the DNA sequence normally associated with most centromeric CENP-A. We did not observe acentric chromosome 3 or evidence of neocentromere formation in our analysis, consistent with the low frequency of such events reported in other systems (Shang et al. 2013; Murillo-Pineda et al. 2021). Together, these findings demonstrate that centromeric islands are dispensable for centromere identity, and that residual CENP-A at flanking sequences is sufficient to support local centromere maintenance following targeted deletion.

A key question arising from the preservation of centromere identity following *Giglio* deletion is whether CENP-A levels at centromere 3 are reduced or maintained. Because the majority of CENP-A normally localizes to the *Giglio* island (Chang et al. 2019), we expected that deletion would substantially decrease CENP-A signal in the absence of compensatory mechanisms. Instead, we observed no significant difference in mean CENP-A intensity between *Giglio*-deleted (G^del^), partially deleted (parG^del^), or intact (G^wt^) centromere 3s. This suggests that total CENP-A levels are maintained following island deletion, consistent with redistribution and/or additional deposition of CENP-A within the flanking satellite sequences.

Although total CENP-A levels were maintained following *Giglio* deletion, the precision of CENP-A organization was not. CENP-A intensity became substantially more variable among centromeres from metaphase spreads containing detectable *Giglio* deletion events, whereas spreads lacking detectable deletions exhibited variability comparable to CR-controls. Thus, deletion of the DNA sequence normally associated with most CENP-A does not impair maintenance of centromere identity or overall CENP-A levels but reduces the fidelity with which the CENP-A domain is re-established.

This reduced precision became even more apparent when we examined CENP-A domain architecture on stretched chromatin fibers. Rather than adopting a uniform architecture following *Giglio* deletion, CENP-A domains exhibited a striking increase in size variability. While some deleted centromeres retained domains similar in size to controls, others expanded significantly, resulting in an approximately tenfold increase in the mean domain size. These findings indicate that *Giglio* deletion does not direct CENP-A toward a new stereotypical architecture but instead removes constraints that normally define the size of the CENP-A domain.

The dramatic increase in domain size variability raised the question of how the expanded CENP-A domain is organized following *Giglio* deletion. In principle, the domain could expand symmetrically into both flanking satellites or preferentially into one side. Our analyses of CENP-A domain size and positioning on extended chromatin fibers showed a directional shift in which Gdel domains were consistently associated with *dodeca*. Importantly, preferential redistribution into *dodeca* was observed only following *Giglio* deletion and not in Gwt centromeres, regardless of genotype, indicating that loss of the island itself rather than CRISPR-mediated DNA damage is associated with CENP-A redistribution.

The preferential redistribution into *dodeca* raises the question of why the two flanking satellites might behave differently. One possibility is that *dodeca* provides a chromatin environment more permissive for CENP-A assembly or retention than *Prodsat*. Alternatively, the residual *Giglio* sequence remaining after deletion may retain directional boundary activity that preferentially restricts spreading toward *Prodsat*. Distinguishing between these possibilities will require targeted manipulation of the flanking satellite sequences.

Collectively, our findings support a model in which centromeric islands serve not simply as preferred substrates for CENP-A assembly, but as structural elements that constrain the size and position of the CENP-A domain. Deletion of *Giglio* releases these constraints, permitting the CENP-A domain to adopt a broad range of configurations while preserving centromere identity. The marked increase in both CENP-A intensity and domain size variability suggests that *Giglio* normally promotes faithful re-establishment of a reproducible centromere architecture following each cell cycle.

An important consideration is that *Drosophila* centromeric islands vary substantially in size and repeat composition, with the centromere 2 island measuring only ∼2 kb compared with the much larger islands found at the other centromeres (Chang et al. 2019). Thus, the constraining function demonstrated here for *Giglio* may not apply uniformly across all *Drosophila* centromeres. Whether smaller or compositionally distinct islands similarly constrain CENP-A domain size and positioning remains an important question for future studies.

This proposed constraining function is conceptually reminiscent of tRNA genes at *S. pombe* centromeres, which are postulated to prevent heterochromatin from spreading into the central core (Scott et al. 2006), and of transitions in DNA methylation demarcated by LINE-1 elements at human centromeres (Altemose et al. 2022; Gershman et al. 2022; Hoyt et al. 2022b). Although these systems differ mechanistically, they suggest that specialized DNA elements can define the permissible boundaries of centromeric chromatin.

More broadly, our findings demonstrate that centromere identity and CENP-A domain organization can be genetically uncoupled. Removal of the DNA sequence normally associated with most centromeric CENP-A does not abolish centromere identity but profoundly alters the architecture of the CENP-A domain. These observations highlight the robustness of epigenetic centromere inheritance while revealing an important role for centromeric DNA in ensuring that this epigenetic state is faithfully re-established with a reproducible chromatin architecture.

Several caveats should be considered when interpreting our findings. First, *Giglio* deletion is mosaic, such that population-based assays, including CUT&Tag, reflect mixtures of cells with and without deletion events, likely underestimating the extent of CENP-A redistribution in fully deleted cells. Second, our analyses were limited to L3 larval brains, and the consequences of *Giglio* deletion may differ in other tissues or developmental stages, particularly in the germ line, where centromere inheritance across generations is most critical. Finally, although CR+ brains exhibited increased mitotic index and apoptosis, we did not detect aneuploidy or overt chromosome segregation defects, suggesting that any effects on kinetochore function are likely to be subtle or are masked by the mosaic nature of the system.

In summary, targeted deletion of the retroelement-rich *Giglio* island demonstrates that centromere identity can be maintained at the native locus despite removal of the DNA sequence associated with most centromeric CENP-A. However, loss of *Giglio* profoundly alters CENP-A domain organization, increasing variability in CENP-A levels, expanding and repositioning the CENP-A domain, and promoting its preferential redistribution into the adjacent *dodeca* satellite (**Fig. 5F**). Together, these findings show that while particular centromeric DNA sequences are not required to maintain centromere identity they may play a critical role in ensuring the faithful re-establishment of CENP-A domain architecture, revealing that the genetic determinants of centromere identity are separable from those that constrain the organization of centromeric chromatin.

## Supporting information

Table S1 - Reagents

## Supplemental figure legends

**Figure S1.**
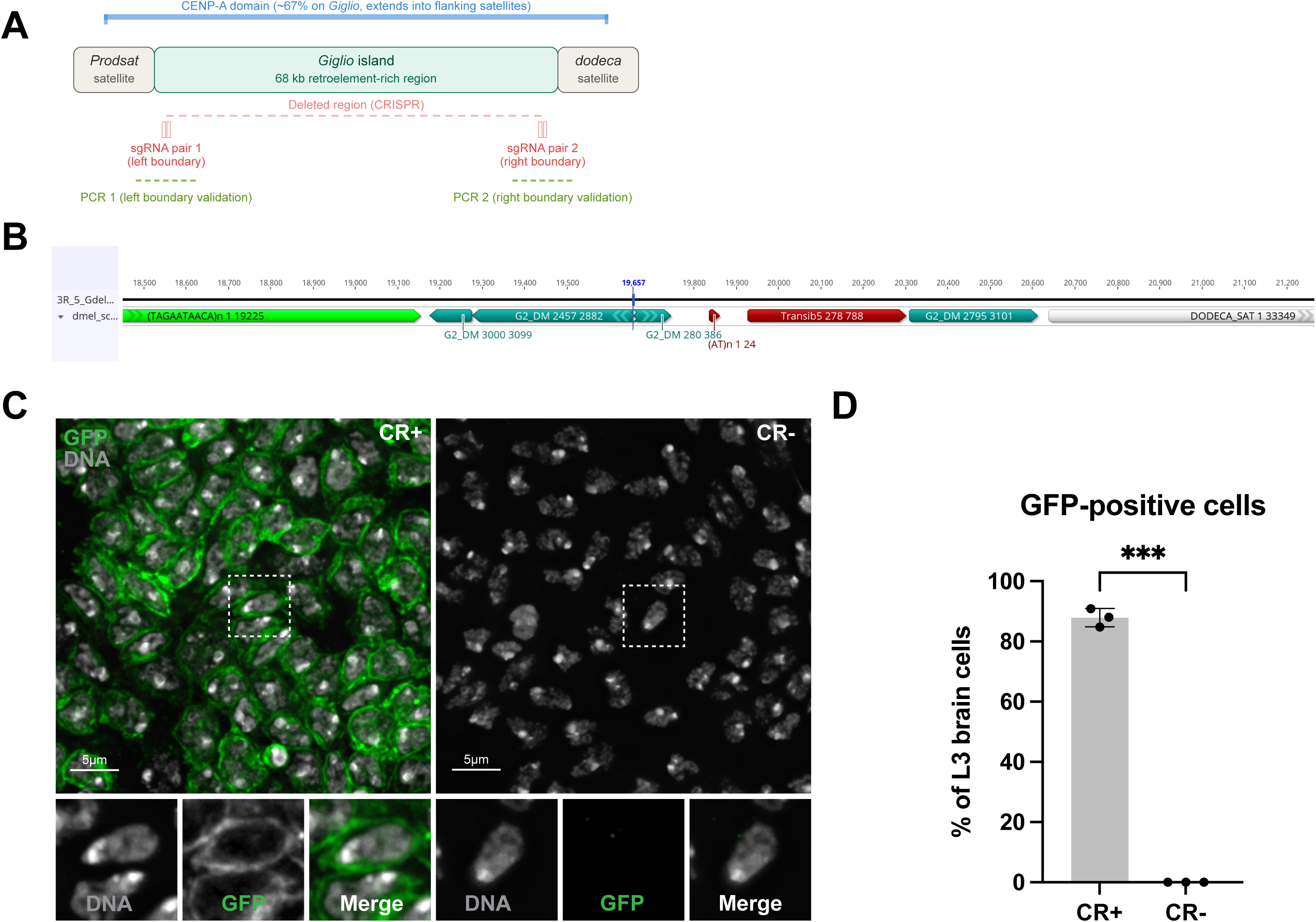
Design of the *Giglio* deletion strategy and validation of GAL4-driven transgene expression in larval brains (related to Fig. 1). A) Schematic (not to scale) of the centromere 3 region showing the *Giglio* island flanked by *Prodsat* and *dodeca* satellite sequences. The CENP-A domain (blue bracket) spans the island, with ∼67% of CENP-A localizing to *Giglio* and the remainder extending into the flanking satellites (Chang et al. 2019). Two pairs of sgRNAs were designed against unique sequences at the left (sgRNA pair 1) and right (sgRNA pair 2) boundaries of the island; the interval predicted to be excised following Fok1:dCas9-mediated cleavage and NHEJ repair is indicated by the red dashed line. Green dashed lines indicate the regions amplified to validate the sgRNA target sequences at each boundary (PCR 1 and PCR 2). B) Repeat annotation of the predicted *Giglio*-deleted (G^del^) allele following excision and NHEJ repair, showing the sequence retained between the *Prodsat* (green) and *dodeca* (gray) satellites. The vertical blue line marks the predicted repair junction. Retained sequences comprise fragments of the *G2/Jockey-3* retroelement (teal), the DNA transposon *Transib5* (red), and a short AT-rich repeat. Numbers beside each feature indicate the coordinates of the aligned segment within the corresponding repeat consensus; coordinates along the top refer to the predicted G^del^ assembly. C) Representative maximum intensity projections of third instar larval brain monolayers from CR+ (left) and CR- (right) animals, showing elav-GAL4-driven UAS-GFP expression (green) and DNA stained with DAPI (gray). Dashed boxes indicate the cells shown at higher magnification below, with the DNA and GFP channels in grayscale alongside the merge. GFP outlines the cortex of each cell. GFP expression in CR+ brains confirms activity of the UAS/GAL4 system and, by extension, expression of the Fok1:dCas9 and sgRNA transgenes; CR- siblings, which lack the GAL4 driver, show no GFP signal. Scale bars, 5 µm. D) Percentage of L3 brain cells expressing GFP in CR+ versus CR- brains. Each point represents one biological replicate (N = 3 brains; n = 600 interphase cells). Bars show the mean, error bars ± SD. ***p < 0.001 (unpaired t-test).

**Figure S2.**
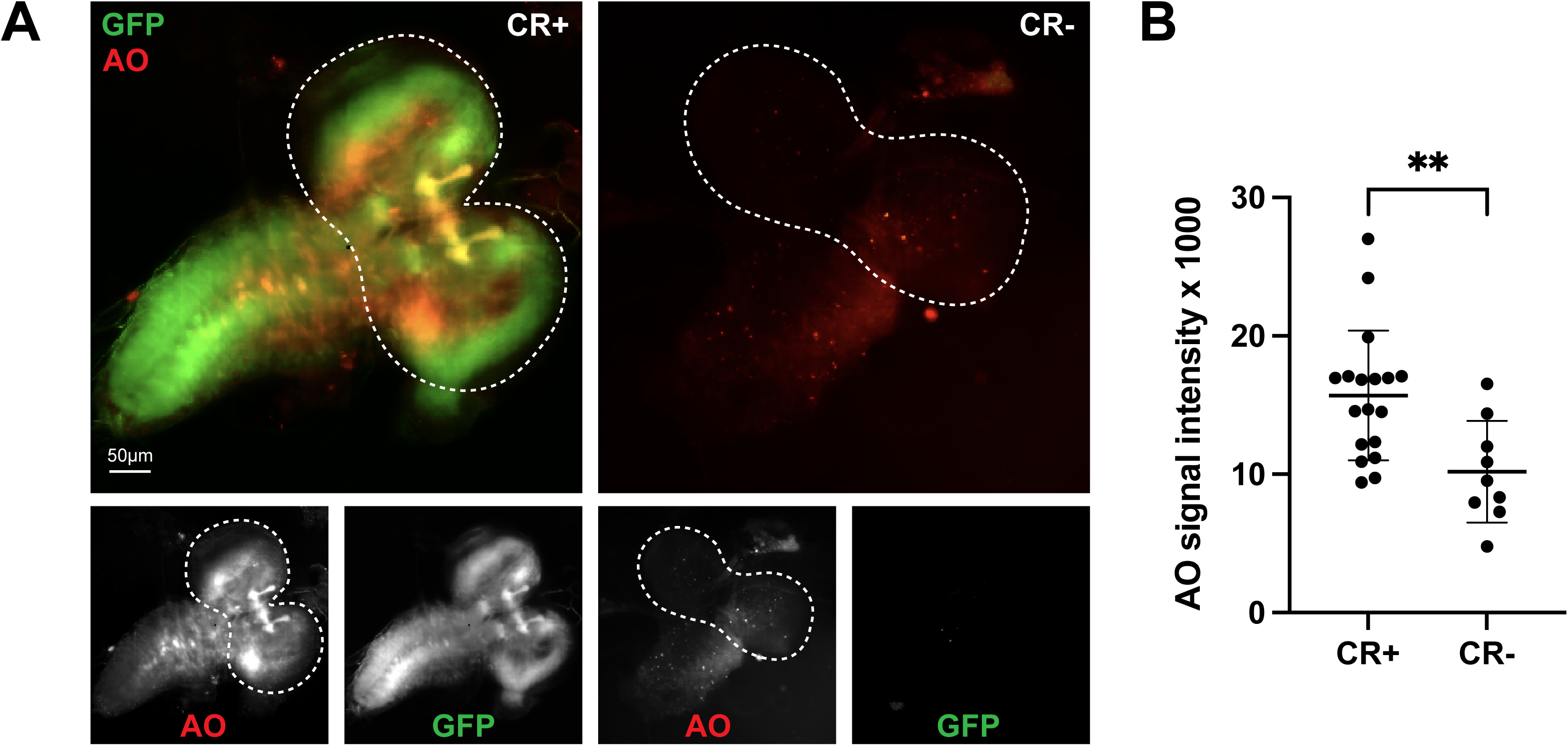
Assessment of cell death in larval brains (related to Fig. 2) A) Representative maximum intensity projections of live L3 brain wholemount from CR+ (left) and CR- (right) animals, stained with acridine orange (AO, red) to detect apoptotic cells. CR+ samples show colocalization of elav>GFP (green) and AO signals, indicating cell death in CRISPR-active neural tissue. CR- brain shows minimal AO staining and lacks GFP fluorescence. A white dashed outline indicates the two optic lobes of the L3 brain. Bottom panels show individual channels (AO, GFP) for each genotype. Scale bar, 50µm. B) AO signal intensity (arbitrary units × 1000) in CR+ versus CR- brains. Each point represents one brain lobe (CR+, n = 9 brains; CR-, n = 5 brains). Lines show the mean ± SD. **p < 0.01 (Welch’s unpaired t-test).

**Figure S3.**
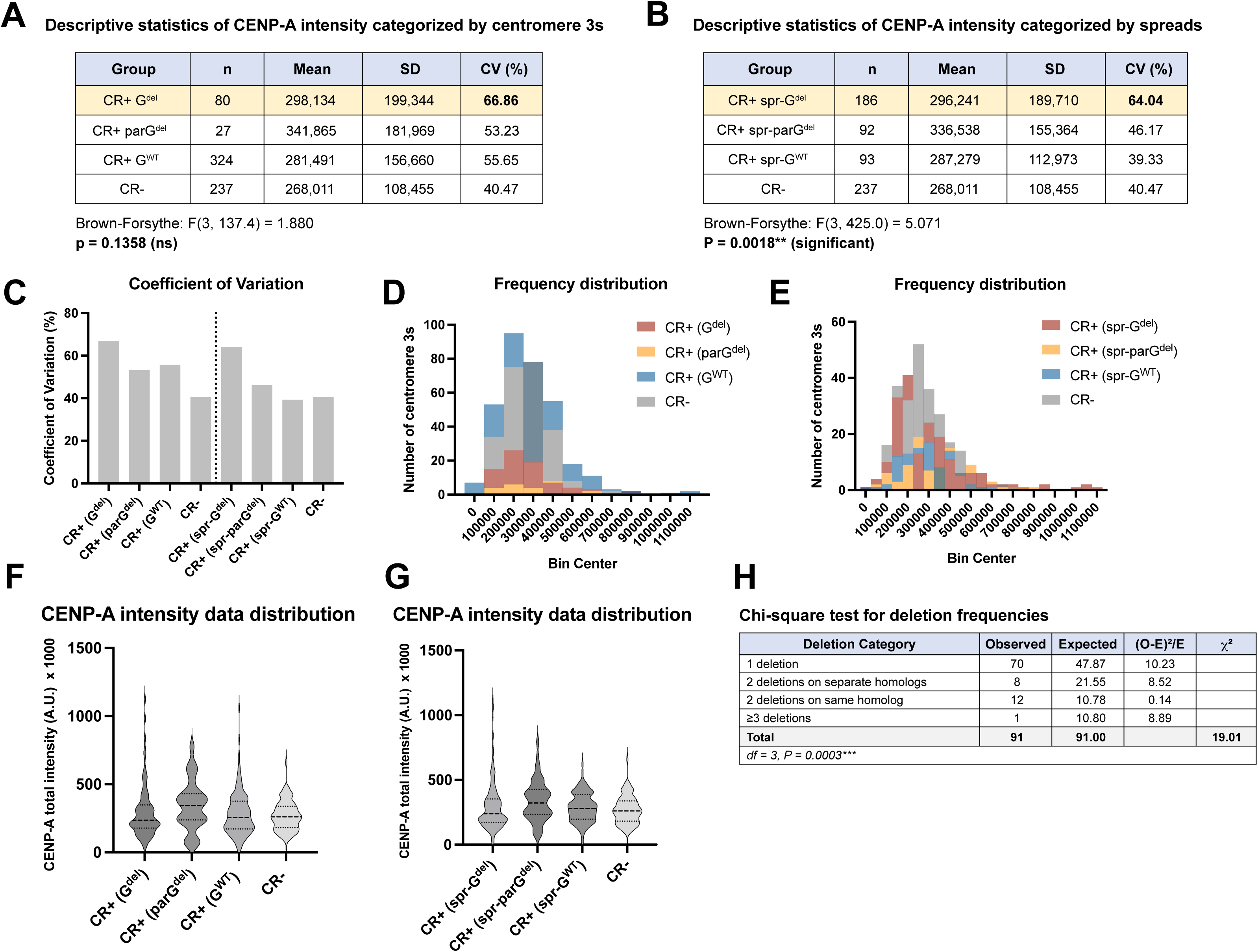
Statistical analysis of CENP-A intensity variance and deletion frequencies (related to Fig. 3) A) Descriptive statistics of CENP-A intensity categorized by centromere 3 genotype. Table showing sample size (n), mean intensity, standard deviation (SD), and coefficient of variation (CV%) for each group: CR+ G^del^ (n = 80 centromere 3s), CR+parG^del^ (n = 27 centromere 3s), CR+ G^wt^ (n = 324 centromere 3s), and CR- (n = 237 centromere 3s). CR+ G^del^ centromeres show the highest coefficient of variation (CV = 66.86%), indicating substantial heterogeneity in CENP-A levels following island deletion. Brown-Forsythe test: F(3, 137.4) = 1.880, p = 0.1358 (ns), indicating no significant difference in variance when categorized by individual centromere genotype. B) Descriptive statistics of CENP-A intensity categorized by cellular genotype (metaphase spread- level classification). Table showing statistics when centromeres are grouped by phenotype of the metaphase spread from which they originated: CR+ spr-G^del^ (cells containing at least one G^del^ centromere 3, n = 186 centromere 3s), CR+ spr-parG^del^ (cells containing at least one par-G^del^ centromere 3, n = 92 centromere 3s), CR+ spr-G^wt^ (CR+ cells with all wild-type centromere 3s, n = 93 centromere 3s), and CR- (n = 237 centromere 3s). CR+ spr-G^del^ show elevated coefficient of variation (CV = 64.04%) compared to CR- controls (CV = 40.47%). Brown-Forsythe test: F(3, 425.0) = 5.071, p = 0.0018 (**), indicating that variance in CENP-A intensity differs significantly when data are categorized by cellular phenotype rather than by individual centromere phenotype. C) CENP-A intensity data distribution by centromere 3 phenotype. Violin plots showing the distribution of CENP-A signal intensity for CR+ spr-G^del^, CR+ parG^del^, CR+ G^wt^, and CR- centromere 3s. Internal markers show the median (solid center line) and quartile boundaries (dotted line). The broader distribution of CR+ G^del^ values is evident compared to other categories. D) CENP-A intensity data distribution by cellular phenotype. Violin plots showing the distribution of CENP-A signal intensity for CR+ spr-G^del^, CR+ spr-parG^del^, CR+ spr-G^wt^, and CR- categories. Internal markers show the median (solid center line) and quartile boundaries (dotted line). CR+ spr-G^del^ shows the broadest distribution, consistent with elevated variance at the cellular level. E) Chi-square test for independence of deletion patterns. Table showing observed versus expected frequencies of deletion categories in two independent experiments. Experiment 1 (CENP-C spreads, n = 150 spreads with at least one deletion) and Experiment 2 (CENP-A spreads, n = 91 spreads with at least one deletion) both show significant deviation from the expected frequencies under a binomial independence model (Experiment 1: χ² = 13.48, df = 4, P = 0.0091**; Experiment 2: χ² = 19.01, df = 3, P = 0.0003***). Observed frequencies show an excess of single deletions and depletion of multi-deletion events, indicating that deletions do not occur independently across the four centromere 3s in a given mitotic cell. Expected frequencies were calculated using deletion probabilities p = 0.51 (Experiment 1) and p = 0.31 (Experiment 2) from the subset of spreads containing at least one deletion.

**Figure S4.**
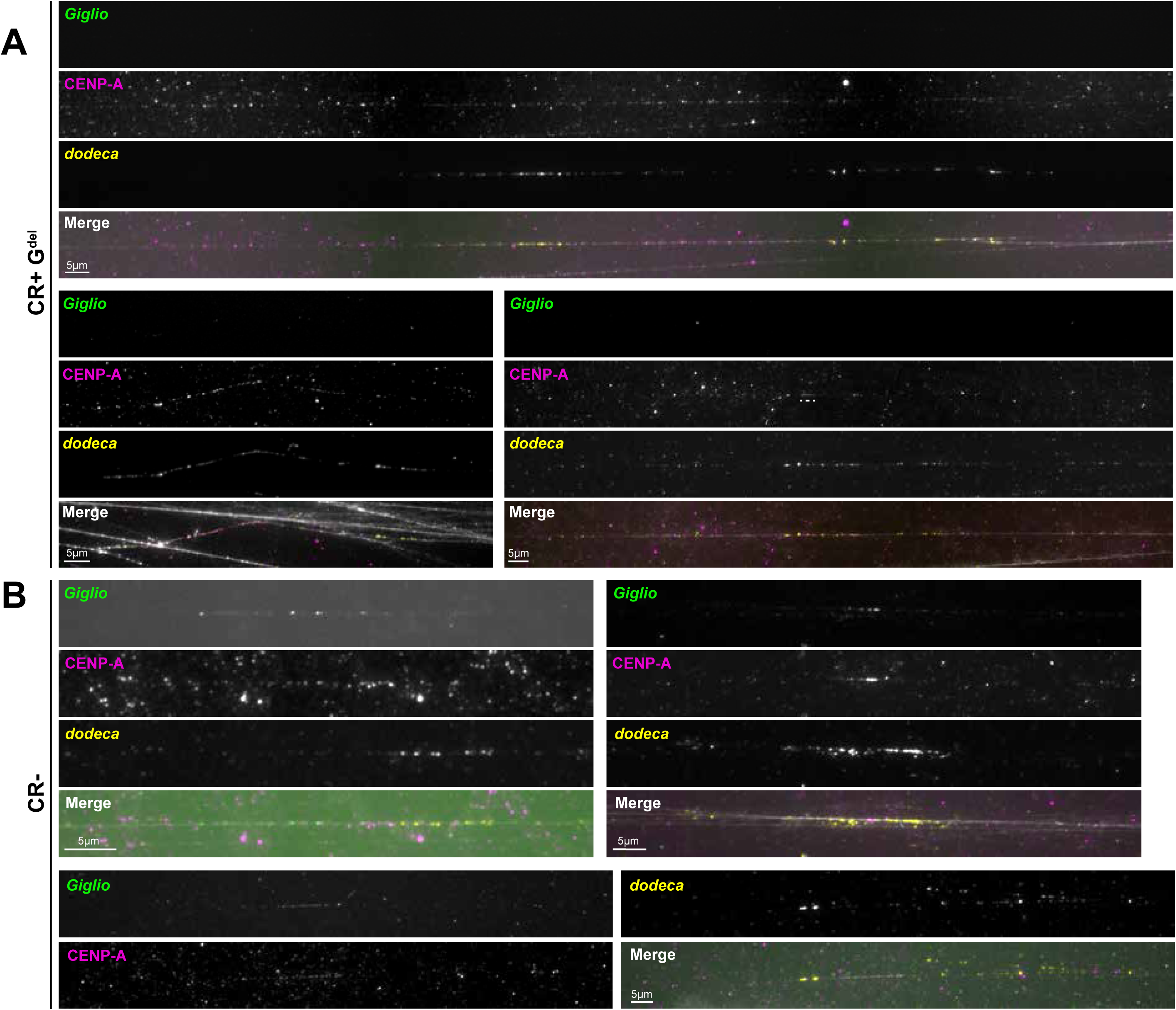
Variability in CENP-A domain size across Giglio-deleted fibers (related to Fig. 4) A) Three additional examples of extended chromatin fibers carrying a G^del^ centromere 3 from CR+ larval brains, illustrating the range of CENP-A domain sizes observed following Giglio deletion. B) Three additional examples of extended chromatin fibers from CR- larval brains. For all fibers, *Giglio* OligoPaint FISH signal is shown in green, CENP-A immunofluorescence in magenta, and the *dodeca* satellite probe in yellow; individual channels are shown in grayscale above the merge. G^del^ fibers lack detectable *Giglio* signal, and the CENP-A domain coincides with the *dodeca*-positive region, whereas CR- fibers retain *Giglio* signal and a shorter CENP-A domain. Fibers shown are representative of those quantified in Fig. 4B-C. Scale bars, 5 µm.

**Figure S5.**
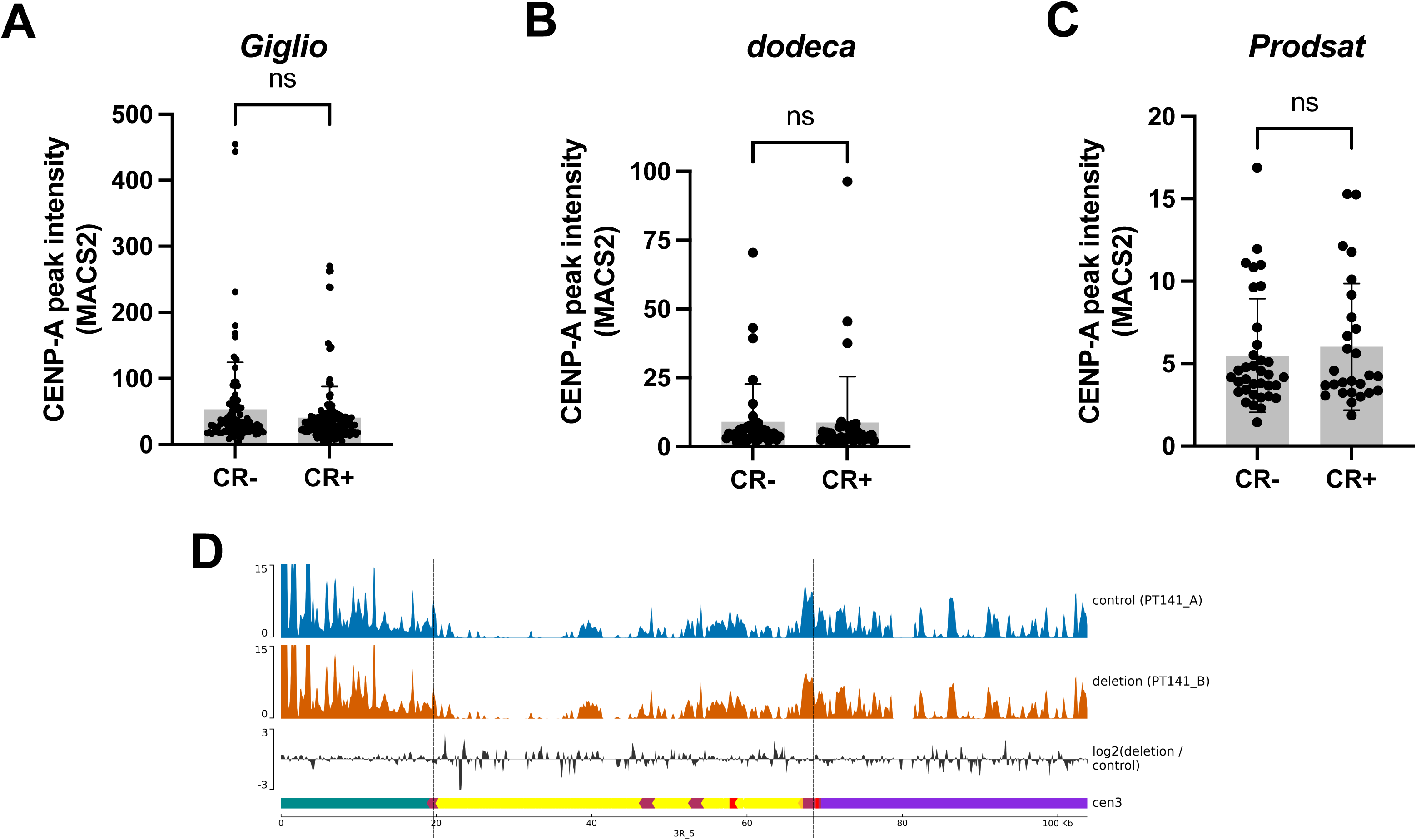
CENP-A peak intensity and whole-genome sequencing coverage across centromere 3 (related to Fig. 5) Intensity of individual CENP-A peaks called by MACS2 across *Giglio* (A), *dodeca* (B), and *Prodsat* (C) in CR- versus CR+ larval brains. Each point represents one called peak, assigned to a region on the basis of at least 50% overlap. Bars show the mean, error bars ± SD. ns = not significant (unpaired t-test). D) Whole-genome sequencing coverage across the centromere 3 contig in CR- and CR+ larval brains. Tracks show read depth for each sample and the log2 ratio of CR+ to CR- coverage. The annotation track indicates *Prodsat* (teal), the *Giglio* island (yellow, with retroelement annotations), and *dodeca* (purple). Vertical dashed lines mark the sgRNA cut sites delimiting the excised interval.

## Tables Legends

**Table S1. Key resources used in this study**

Reagents, equipment, software, probes, fly stocks, plasmids, oligonucleotides, and antibodies used in this study. The ‘*Reagents’* sheet is organized by category: chemicals, enzymes, and commercial kits (supplier and catalog number); equipment (manufacturer and model); software (developer and version or RRID); DNA and FISH probes (probe type, source, and sequence where applicable); *Drosophila* stocks (short alias, full genotype, and source or Bloomington *Drosophila* Stock Center identifier); plasmids and constructs (source and RRID or accession); oligonucleotides and primers (application and sequence); and primary and secondary antibodies (supplier, catalog number, and working dilution). Stocks and plasmids generated in this study are indicated as “This paper.” The ‘FokIdCas9_sequences’ sheet gives the nucleotide sequences of the two synthetic fragments used to assemble Fok1:dCas9 (2,420 bp each, sharing a 30 bp overlap for Gibson assembly) and of the complete pUASz-Fok1:dCas9 construct (12,485 bp).

**Table S2. sgRNA sequences, target sites, and off-target analysis**

Off-target analysis of the four sgRNAs used to excise the *Giglio* island, performed with BLASTN v2.7.1 against the heterochromatin-enriched *D. melanogaster* genome assembly (Chang et al., 2019). The ‘Off_target_analysis’ sheet lists perfect and single-mismatch hits only; for each, it gives the contig on which a match was found, aligned sequence, the number of mismatches relative to the sgRNA, the coordinates of the match, and whether the site corresponds to the intended target. The ‘gRNA_summary’ sheet gives, for each gRNA, its sequence, the boundary targeted, and the number of sites in the assembly matching with zero, one, or two mismatches, together with the number of perfect-match off-target sites (excluding the intended on-target site).

## Methods

### Drosophila stocks and handling

All *Drosophila melanogaster* stocks (**Table S1**) were reared on standard cornmeal-molasses-yeast medium (https://bdsc.indiana.edu) at 25°C. GAL4 driver lines were crossed with w[*]; P{w[+mC]=UASt-4sgRNAs}attP64/TM3, Sb to generate GAL4 driver; UAS-sgRNAs flies. To ensure that the centromere 3s under analysis originated from the characterized iso-1 reference stock, iso-1 chromosome 3s were introgressed by crossing y^1^ w^67c23^; P{w[+mC]=UASz-FokI:dCas9}attP40/CyO; D/TM3,Tb flies to iso-1 and selecting against D and TM3. All stocks were maintained as heterozygous stocks, and homozygotes from UAS-FokI:dCas9/CyO; +^iso-1^ were selected for setting up crosses (**Fig. 1B**).

### Guide RNA design, plasmid cloning, and transgenic fly generation

The sgRNA transgenic stock was generated by φC31-mediated integration of pCFD6-UASt:4sgRNAs at the attP64 landing site on chromosome 3 after injection into y^1^ w^67c23^; P{CaryP}attP64 embryos by a commercial service (BestGene). The four sgRNAs were designed against sequences near both the ends of *Giglio* island using the “Find CRISPR sites” function in Geneious. Two compatible pairs of sgRNAs located 13-15 bp apart with high specificity scores based on off-target analysis against the *D. melanogaster* heterochromatin-enriched genome assembly (Chang et al., 2019) were selected (**Fig. S1C**) and the presence of the target genomic sequence was verified by PCR followed by Sanger sequencing in the w[*]; P{w[+mC]=UASt-4sgRNAs}attP64 stock. The four sgRNAs were: 5’-AAATACAGAAGGCTGACTTG-3’, 5’-GACGGCGACTACATCCCTAC-3’, 5’-CCAAAGAACTAAGTGCGATG-3’, and 5’-CCTCTGCTCTGCTCCTCTGC-3’.

Off-target alignments were verified using BLAST against the same genome assembly (Chang et al., 2019; **Table S2**). The pCFD6-UASt:4sgRNAs plasmid was generated from pCFD6 (gift from Simon Bullock) following the protocol described in (Port and Bullock 2016) (crisprflydesign.org). To clone the four sgRNAs, three fragments were generated using PCR primers listed in **Table S1**. The pCFD6 plasmid was linearized by restriction digestion with BbsI-HF (NEB), and the amplified sgRNA fragments were assembled into the linearized vector using NEBuilder HiFi DNA Assembly. The final construct was verified by Sanger sequencing.

The UAS-Fok1:dCas9 line was generated by φC31-mediated integration of pUASz-Fok1:dCas9 at the attP40 landing site on chromosome 2 after injection into y^1^ w^67c23^; P{CaryP}attP40 embryos by a commercial service (BestGene). The pUASz1.0 plasmid (DGRC) was linearized using KpnI-HF (NEB). The Fok1:dCas9 transgene, based on pAct-Fok1:dCas9 (gift from Simon Bullock), was synthesized as two fragments by Azenta Life Sciences to accommodate size constraints for gene synthesis. The first fragment spanned the Fok1 nuclease and part of the dCas9 coding sequence, and the second fragment spanned the remaining dCas9 sequence, with 20 bp overlapping regions for Gibson assembly (**Table S1**). These fragments were assembled into the linearized pUASz1.0 backbone plasmid using NEBuilder HiFi DNA Assembly. The final construct was verified using whole plasmid sequencing (Eurofins Genomics).

### Genetic crosses and GAL4/UAS expression

Expression of the CRISPR system was under the control of GAL4 drivers. Crosses used UAS-sgRNA males and UAS-Fok1:dCas9 virgin females.

For our analyses in mitotic cells, interphase cells, chromatin fibers, and for our CUT&Tag, the neuronal elav-GAL4 was used to drive expression in the larval brain. The larval progeny were genotyped based on GFP fluorescence in larval brains, where GFP-positive brains were designated as CR+ and GFP-negative brains as CR-controls.

For expression in the developing eye, eyeless-GAL4 driver was used. eyeless-GAL4/CyO; UAS-sgRNAs flies were crossed with UAS-Fok1:dCas9, and the hatched adult progeny were segregated based on wing phenotype: straight-winged flies were CR+, curly-winged flies were CR- or control. Adults were anesthetized with CO_2_, screened, and imaged. Both genotypes were scored for changes in the eye shape and damage to the ommatidia.

For the viability assay, actin-GAL4 was used to drive ubiquitous expression. Three to five actin-GAL4/CyO; UAS-sgRNAs males were crossed with a single UAS-Fok1:dCas9 virgin female. Ten such crosses were established at 25°C. Simultaneously, a similar setup was done for each of the parental lines to rule out genotype-related viability defects. Flies were allowed to mate and lay eggs for 72 h before being removed from the vials. Eclosed progeny were counted and removed from the vials daily.

### Metaphase spreads preparation from larval brains

Brains from third instar larvae were dissected in 0.7% NaCl, and all attached tissue and mouth parts were removed with forceps. Brains were immersed in 0.5% sodium citrate solution for 8 min in a spot well dish. Each brain was transferred to a 4-µl drop of fixative (45% acetic acid, 2% formaldehyde) on a siliconized coverslip and fixed for 6 min. Three brains per coverslip were used. A poly-lysine-coated glass slide was inverted onto the coverslip to create a sandwich. The slide was flipped, placed between sheets of bibulous paper, and gently pressed to remove excess fixative. Brains were squashed by pressing firmly on the coverslip with the thumb. Slides were immersed in liquid nitrogen until bubbling ceased, and coverslips were removed with a razor blade. Slides were rehydrated in PBS for 5 min before proceeding with IF-FISH.

### Acid-free metaphase spreads

Brains from third instar larvae were dissected in 0.7% NaCl. Brains were transferred to 1× colchicine and incubated for 1 h at 25°C. Brains were then transferred to 0.5% sodium citrate solution and incubated for 8 min. Each brain was transferred to a 5-µl drop of 4% formaldehyde on a siliconized coverslip, immediately squashed gently between the slide and coverslip using a pencil eraser tip, and then compressed with a hand clamp. Slides remained clamped for a total of 5 min from the start of squashing. Slides were then flash-frozen in liquid nitrogen until bubbling ceased, and coverslips were removed with a razor blade. Slides were rehydrated in 1× PBS and processed immediately for IF-FISH.

### IF-FISH

For IF-FISH, all the washes at room temperature were done on a shaker. Slides were transferred to PBST (PBS containing 0.1% Triton X-100) for 10 min, then blocked in 1% BSA in PBST for 30 min. Primary antibodies, anti-CENP-C (Guinea pig, 1:500) or anti-CENP-A (Chicken, 1:500), were diluted in blocking solution, applied to slides, and incubated for 1 h at room temperature or overnight at 4°C. Slides were washed three times for 5 min each in PBST. Secondary antibodies, anti-guinea pig Alexa Fluor 546 (Goat, 1:500) or anti-chicken Alexa Fluor 546 (Goat, 1:500), were diluted in blocking solution, applied to slides, and incubated for 45 min at room temperature. Slides were washed three times for 5 min each in PBST. For FISH, slides were washed twice for 5 min each in 2× SSCT (2× SSC containing 0.1% Tween-20), then washed in 2× SSCT containing 50% formamide for 10 min at 37°C. Probe mix was prepared containing 6.25 µl of 4× hybridization buffer (8× SSC, 0.4% Tween-20, 40% dextran sulfate), 40 pmol of dodeca LNA satellite probe (Chang et al. 2019), 10 pmol of *Giglio* Oligopaint probe (Chang et al. 2019), 0.1 µl of Alexa-488-labeled secondary oligo (100 µM), 1 µl of RNase cocktail (Invitrogen), and 50% formamide, adjusted to 25 µl with ultrapure water. Probe mix (25 µl) was applied to each slide, covered with a coverslip, and sealed with rubber cement. Slides were denatured at 85°C for 10 min, then incubated overnight at 42°C in a dark, humid chamber. Rubber cement was removed, and slides were washed in 2× SSCT for 10 min at 60°C, followed by two washes for 15 min each in 2× SSCT at room temperature, and one wash for 5 min in PBS. Slides were mounted in SlowFade Gold containing DAPI (1.5 µg/ml).

### Larval brain monolayer preparations and immunofluorescence

Third instar larvae were collected and washed twice in PBS to remove food debris. Larvae were dissected, and heads were transferred to a well containing PBS. The tissues were fixed in 3.7% formaldehyde in PBS for 5 min with rotation, then washed four times for 5 min each in PBS. The tissues were transferred to dissociating solution (0.04% Dispase/Collagenase in PBS) and incubated at 37°C for 10 min. Tissues were transferred to a new well containing PBS, and brains were dissected. Brains were transferred to 2-µl drops of PBS on siliconized coverslips, with up to four drops per coverslip. Tissues were squashed onto a glass slide and immediately flash-frozen in liquid nitrogen. Coverslips were removed with a razor blade, and 400 µl of fixative (3.7% formaldehyde in PBS) was immediately applied to the squashed tissue within 30 sec. Slides were fixed for 5 min, then washed four times for 5 min each in PBS.

Immunofluorescence on dissociated larval tissues was performed as described previously (Blower and Karpen 2001; Peng and Karpen 2009) with modifications. Slides were blocked in blocking solution (1% milk in PBS, 0.1% Triton X-100) for 1 h with rotation at room temperature. Primary antibodies anti-H2AvD-pS137 (referred to as γH2Av) antibody (rabbit,1:250; Rockland), anti-phosphorylated H3 serine 10 (rabbit, 1:1000; Sigma Aldrich), and anti-CENP-A (chicken; **Table S1**) were diluted in blocking solution, applied to slides, and incubated overnight at 4°C in a humid chamber. Slides were washed three times for 5 min each in blocking solution. Secondary antibodies, anti-rabbit Alexa Fluor 488 (Goat, 1:500), and anti-chicken Alexa Fluor 546 (**Table S1**) were diluted in blocking solution, applied to slides, and incubated for 45 min at room temperature in a dark, humid chamber. Slides were washed three times for 5 min each in PBS and mounted in SlowFade Gold containing DAPI (Thermo Fisher).

### FISH-IF on chromatin fibers

Chromatin fibers (Sullivan 2010) were prepared from third instar larval brains. All wash and incubation steps were performed without agitation to preserve fiber integrity and ensure consistent IF-FISH staining. Three brains per slide were dissected in 0.7% NaCl and transferred within 15 min to a multi-well plate containing 250 µl of 0.5% (w/v) sodium citrate. Brains were incubated at 37°C for 10 min.

Dispase/Collagenase solution (250 µl; 40 µg in Schneider’s medium; Sigma) was added to the sodium citrate solution and mixed by pipetting. Brains were incubated at 37°C for 10 min, and the cell suspension was transferred to a single-chamber cytofunnel and centrifuged onto poly-L-lysine-coated glass slides at 1200 rpm (90 × g) for 5 min using a Cytospin 3 centrifuge (Shandon). Slides were immediately placed in lysis buffer (500 mM NaCl, 25 mM Tris-HCl pH 7.5, 250 mM Urea, 1% Triton X-100) for 16 min without agitation. Slides were removed from the lysis buffer by allowing the buffer to drain, then immediately immersed in fixative (4% formaldehyde in PBS) without agitation. Fixative was drained, and slides were immersed in PBS for 2 min, then transferred to PBS containing 0.1% Triton X-100 for 10 min. Slides were washed three times for 5 min each in PBS, then once in 2× SSC containing 0.1% Tween- 20 (SSCT) for 10 min.

For FISH, slides were washed in 2× SSCT containing 50% formamide for 10 min at 37°C. Probe mix was prepared containing 6.25 µl of 4× hybridization buffer (8× SSC, 0.4% Tween-20, 40% dextran sulfate), 40 pmol of *dodeca* LNA satellite probe (Chang et al., 2019), 10 pmol of *Giglio* Oligopaint probe (Chang et al., 2019), 0.1 µl of Alexa-488-Sec6 secondary oligo 100 µM; **Table S1**), 1 µl of RNase cocktail (Invitrogen), and 50% formamide, adjusted to 25 µl with ultrapure water. Slides were briefly drained, and 25 µl of probe mix was applied per slide and covered with a HybriSlip coverslip sealed with rubber cement. Slides were incubated overnight at 42°C in a humid chamber. HybriSlips were removed with a razor blade, and the slides were washed in 2× SSCT for 10 min at 60°C, followed by 2× SSCT for 15 min at room temperature, and 0.2× SSC for 10 min at room temperature.

For IF, slides were washed three times for 5 min each in PBS, and the slides were blocked in 1.5% BSA in PBS for 30 min without agitation. Primary antibody anti-CENP-A (chicken, 1:100; **Table S1**) was diluted in blocking solution, applied to slides (50 µl per slide), covered with parafilm, and incubated overnight at 4°C in a humid chamber. Slides were washed three times for 5 min each in PBS without agitation.

Secondary antibody, anti-rabbit Alexa Fluor 546 (Goat, 1:500), was diluted in blocking solution, applied to slides (50 µl per slide), and incubated for 1 h at 25°C. Slides were washed three times for 5 min each in PBS. Slides were mounted in SlowFade Gold with DAPI (1.5 µg/ml) (Thermo Fisher) under 22 × 40-mm coverslips.

### Imaging

Images of mitotic spreads, monolayers, and stretched chromatin fibers were acquired at 25°C using an inverted DeltaVision Ultra microscope (Leica Microsystems) equipped with a sCMOS pco.edge detector camera. All the slides were imaged using a 100×/1.40 NA oil objective (Olympus) with 0.2µm z-stacks for spreads and monolayers and 0.1µm z-stacks for stretched chromatin fibers. Very long chromatin fibers were imaged using the panels option in Acquire Ultra software and stitched together without deconvolution. Whole-mount tissues were imaged using a 40×/1.35 NA oil objective (Olympus) in panels and stitched together. Image acquisition was performed using DeltaVision Ultra Image Acquisition software or softWoRx software (Applied Precision). Images acquired with 40× or 100× oil objectives were deconvolved for five cycles using the conservative algorithm in softWoRx. Maximum intensity projections were generated from in-focus z-slices. A uniform intensity scale was applied to all images prior to export.

All images were minimally adjusted in Photoshop (Adobe) and assembled into figures using Illustrator (Adobe).

### *Giglio* deletion quantification in metaphase spreads

Chromosome 3 centromeres were identified in metaphase spreads using *dodeca* satellite FISH signal. Colocalization of *Giglio* OligoPaint FISH signal with CENP-C or CENP-A immunofluorescence signal was assessed visually. Centromeres displaying *Giglio* signal were categorized as G^wt^, centromeres with very faint and/or small *Giglio* signal were categorized as parG^del^, and centromeres lacking *Giglio* signal, even after increasing the brightness of the signal, were categorized as G^del^.

### Colocalization quantification for γH2Av at centromeres

Centromeres were identified in interphase cells using the CENP-A immunofluorescence signal. Colocalization of γH2Av IF signal with centromeres was assessed visually and scored as positive if the signal overlapped with CENP-A signal.

### Mitotic index quantification in larval brain monolayers

Nuclei/chromosomes positive for phosphorylated histone H3 serine 10 (pH3S10) IF were scored as in mitosis. Total cells per field were counted using DAPI, regardless of pH3S10 status. Mitotic index was calculated as the percentage of pH3S10-positive cells relative to total. Mitotic indices were quantified for each brain replicate per genotype and plotted and analyzed using GraphPad Prism.

### Cell death assay in larval brains

Cell death was assessed using acridine orange (AO) staining, adapted from Hong et al. (2012). Brains from early third instar larvae were dissected in PBS and incubated in 0.2 µM AO solution (Sigma-Aldrich, A6014-10G; diluted from stock in PBS) for 5 min. Brains were rinsed in PBS for 1 min, mounted in halocarbon oil, and imaged within 10 min of staining completion using an inverted DeltaVision Ultra microscope (Leica Microsystems). Seven to ten brains per genotype were analyzed for cell death. Brain lobes displaying a prominent AO banding pattern indicative of endogenous developmental apoptosis were excluded from the analysis to isolate experimentally induced cell death.

### Fluorescence CENP-A intensity quantification

To quantify CENP-A signal intensity at centromeres in metaphase spreads, non-deconvolved in-focus z-slices were combined as maximum intensity projections using softWoRx. Polygons were drawn around each centromere 3 CENP-A focus and three nearby background regions using the edit polygons tool.

Total fluorescence intensity and area were measured for each polygon. CENP-A signal intensity was calculated by subtracting the mean background total intensity per unit area from the centromeric total intensity per unit area. Signal intensities were plotted and analyzed using GraphPad Prism. Similar steps were performed to quantify CENP-A signal of centromeres X, Y, 2, and 4.

To quantify AO staining for assessing cell death, non-deconvolved in-focus z-slices were combined as maximum intensity projections using softWoRx. Polygons were drawn around each lobe of the larval brain and three nearby background regions using the edit polygons tool. Total fluorescence intensity and area were measured for each polygon. AO signal intensity was calculated by subtracting the mean background total intensity per unit area from the AO total intensity per unit area. Signal intensities were plotted and analyzed using GraphPad Prism.

### Analyzing centromere 3 on stretched chromatin fibers

Well-stretched, single, and isolated chromatin fibers were selected for analysis of *Giglio* deletion and CENP-A domain position at centromere 3. Fibers displaying CENP-A immunofluorescence signal colocalizing predominantly with *Giglio* Oligopaint FISH signal, with some extension into adjacent dodeca-positive regions and/or unstained flanking regions, were categorized as G^wt^. Fibers displaying CENP-A immunofluorescence signal colocalizing predominantly with *dodeca* satellite FISH signal, with no detectable *Giglio* signal within the CENP-A domain or adjacent dodeca-positive region, were categorized as G^del^.

### Measuring CENP-A domain and *Giglio* island using stretched chromatin fibers

Maximum intensity projections of well-stretched, single, and isolated chromatin fibers were used to measure CENP-A domain length in G^wt^ and G^del^ fibers. A line was drawn across the entire length of each CENP-A domain using the line tool in ImageJ, and line length was measured in µm using the measure function. CENP-A domain lengths were plotted and analyzed using GraphPad Prism. *Giglio* island length was measured using the same method and plotted separately.

### Quantification of CENP-A overlap with *dodeca* in stretched chromatin fibers

The same maximum intensity projections used to measure CENP-A domain length were used to quantify the extent of CENP-A overlap with the *dodeca* satellite in G^wt^ and G^del^ fibers from CR+ brains. For each fiber, a line was drawn along the portion of the CENP-A domain coinciding with *dodeca* FISH signal using the line tool in ImageJ, and its length was measured in µm using the measure function. The total CENP-A domain length for the same fiber was measured as described above and recorded alongside this value. The proportion of the CENP-A domain overlapping *dodeca* was then calculated for each fiber as the length of the *dodeca*-coincident segment divided by the total CENP-A domain length, expressed as a percentage. These values were plotted and analyzed using GraphPad Prism.

### CUT&Tag in larval brains

CUT&Tag was performed on third instar larval brains following the published protocol with modifications (Kaya-Okur et al. 2019). Concanavalin A-conjugated magnetic beads (EpiCypher) were resuspended, and 3.4 µl of beads per sample were transferred to a PCR tube, placed on a magnetic stand, and the liquid was replaced with cold bead activation buffer (20 mM HEPES pH 7.9, 10 mM KCl, 1 mM CaCl₂, 1 mM MnCl₂) at 20 µl per sample and incubated for 10 min at room temperature. Third-instar larvae were washed in PBST, and 15 or 20 brains per sample were dissected in PBS within 10 min and transferred to a chilled tube containing 200 µl of ice-cold, EDTA-free nuclei extraction (NE) buffer (20mM HEPES pH 7.9, 150mM NaCl, 1× Roche cOmplete EDTA-free protease inhibitor, 0.1% Triton X-100, and 20% glycerol), followed by a brief spin. Nuclei were released by slow passage through a 30-gauge needle 5 - 10 times. Heavy debris was cleared by centrifugation at 100 × g for 1 min at 4°C, and the supernatant was transferred to a fresh chilled tube. Nuclei were pelleted at 1300 × g for 5 min at 4°C, and the supernatant was removed. Nuclei were resuspended by gentle pipetting in Wash150+ buffer (20mM HEPES pH 7.9, 150mM NaCl, 1× Roche cOmplete EDTA-free protease inhibitor, 0.05% Triton-X 100) at a volume of 40 µl per reaction. Aliquots of 40 µl of this nuclei suspension were added to each sample tube containing 20µl of activated beads, mixed by flicking, and incubated for 5 min at 25°C. Tubes were placed on a magnetic stand, supernatant was removed, and beads were resuspended in 20 µl of primary antibody dilution buffer (1:100 in Wash150+ supplemented with 0.5 mM spermidine, 1× Roche cOmplete EDTA-free protease inhibitor, 2 mM EDTA, and 0.12% BSA). Primary antibodies used were rabbit anti-CID (Active Motif) for CENP-A profiling, rabbit anti-H3K27me3 (Cell Signaling) as a positive control, and rabbit IgG (EpiCypher) as a negative control, all at 1:100 dilutions. Samples were incubated overnight at 4°C.

On day two, tubes were placed on a magnetic stand, supernatant was removed, and beads were resuspended in 20 µl of secondary antibody dilution (guinea pig anti-rabbit, 1:100 in antibody dilution buffer). Samples were incubated for 1 h at 25°C. Beads were captured on a magnetic stand, supernatant was removed, and beads were resuspended in 20 µl of pAG-Tn5 solution (CUTANA pAG-Tn5; EpiCypher; 1 µl per 20 µl Wash300+ (20 mM HEPES pH 7.5, 300 mM NaCl, 0.05% Triton X-100, 0.5 mM spermidine, 1× protease inhibitor) and incubated for 1 h at 25°C without agitation. Beads were washed once with Wash300+ buffer. Tagmentation was initiated by resuspending beads in 20 µl of tagmentation buffer (Wash300+ buffer containing 10 mM MgCl₂) followed by incubation for 1 h at 25°C. Beads were washed with 10 mM TAPS pH 8.5, resuspended in 5 µl of SDS-TAPS release buffer (10 mM TAPS pH 8.5, 0.1% SDS, thermolabile Proteinase K; NEB). Samples were incubated for 1 h at 37°C followed by enzyme inactivation at 55°C for 15 min.

For library amplification, 15 µl of 0.67% Triton X-100 was added to quench SDS, followed by 2 µl each of universal i5 primer and a uniquely barcoded i7 primers (Buenrostro et al. 2015). CUTANA High-Fidelity 2× PCR Master Mix (25 µl; EpiCypher) was added to each sample. PCR was performed as follows: 58°C for 5 min, 72°C for 10 min, 98°C for 30 s, then 14 cycles of 98°C for 10 s and 60°C for 10 s, followed by 72°C for 1 min and a hold at 8°C. Libraries were purified using AMPure XP magnetic beads (Beckman Coulter) at a 1.3:1 ratio. Beads were incubated with samples for 10 min, captured on a magnetic stand for 5 min, washed once with 500 µl of 80% ethanol for 30 sec, and eluted in 25 µl of 10 mM Tris-HCl pH 8.0. Library size distribution and concentration were assessed using an Agilent 4200 TapeStation with D1000 reagents. Libraries were pooled at equimolar concentrations and sequenced using 150-bp paired-end mode on the Illumina Nova seq X plus. We obtained between 62.5-196.8 million mapped reads per library.

### CUT&Tag data processing and analysis

Read quality was assessed using FastQC, and adapters were trimmed using Trim Galore with the following parameters: quality threshold of 20, Nextera adapter removal, minimum length of 35 bp, and Phred33 quality encoding. Trimmed paired-end reads were aligned to a custom *D. melanogaster* genome assembly containing heterochromatin-enriched scaffold sequences (Chang et al. 2019) using Bowtie2 with default parameters. Alignments were converted to BAM format, sorted, and indexed using SAMtools.

Because the IgG control library was pooled at approximately 25-fold higher concentration than the target libraries and was correspondingly over-sequenced, IgG alignments were downsampled to 4% of reads using SAMtools view with a fixed seed, restoring a sequencing depth comparable to that of the CENP-A libraries. The downsampled IgG alignments were used as the control for all subsequent peak calling.

For peak calling, BAM files were converted to BED format using BEDTools, genome coverage was calculated to generate bedGraph files, and these were converted to bigWig format using bedGraphToBigWig (UCSC). Peaks were called using MACS2 with paired-end BAM input (-f BAMPE), the *Drosophila* effective genome size (-g dm), a q-value threshold of 0.01, bedGraph output (-B), and summit calling enabled (--call-summits), using the downsampled IgG BAM as the control. Fraction of reads in peaks (FRiP) scores were calculated by determining the proportion of mapped reads overlapping MACS2 peak regions using BEDTools intersect.

For visualization, read coverage was normalized to counts per million (CPM) using bamCoverage from deepTools with a bin size of 10 bp, read extension to the full fragment length, and read centering. Log2 fold-change tracks were generated from CPM-normalized bigWig files using bigwigCompare from deepTools with a pseudocount of 1, skipping bins with zero coverage in both samples. Signal matrices were computed over regions of interest using computeMatrix scale-regions, and profile plots were generated using plotProfile from deepTools.

### Whole Genome Sequencing and analysis

Genomic DNA was isolated from 20 L3 larval brains per genotype using the DNeasy Blood & Tissue kit (Qiagen). Libraries were generated with the NEBNext Ultra II FS DNA Library Prep kit (NEB) and sequenced in 150-bp paired-end mode on the Illumina NovaSeq X Plus.

Adapters were detected and trimmed using fastp with overlap-based adapter detection for paired-end reads and a minimum retained read length of 50bp. Read quality was assessed using FastQC before and after trimming. Trimmed reads were aligned using Bowtie 2 to the same heterochromatin-enriched *D. melanogaster* assembly used for CUT&Tag (Chang et al. 2019), with the --sensitive preset and a maximum concordant fragment length of 1000 bp. Aligned BAM files were then indexed.

Coverage tracks were generated using bamCoverage from deepTools with a bin size of 50 bp, read extension to the full fragment length, exclusion of marked duplicates, and CPM normalization. Relative coverage between the genotypes was calculated as Log2(CR+/CR-) using bamCompare on uniquely mapped reads (MAPQ≥20).

## Data availability

All code for analyses and figures is available on https://github.com/MelloneLab/gigliodeletion/releases/tag/v1.0.0

All sequencing data is available on NCBI under Bioproject PRJNA1515322.

## Authors contributions

Conceptualization, methodology, resources, writing –original draft, writing –review and editing, visualization, and project administration: P.P. and B.G.M. Validation, and formal analysis: P.P. Investigation: P.P. and D.T. (apoptotic assay). Supervision and funding acquisition: B.G.M.

## Acknowledgements

This work was funded by NIH award R35GM131868 to B.G.M. We thank Bo Reese and the University of Connecticut Center for Genome Innovation in the Institute for Systems Genomics for assistance with sequencing, Noah Reid and Adam Rork (Computational Biology Core, Institute for Systems Genomics), and the University of Connecticut High-Performance Computing for computational support. Plasmids from the Drosophila Genomics Resource Center (supported by NIH grant 2P40OD010949) and stocks obtained from the Bloomington Drosophila Stock Center (NIH P40OD018537) were used in this study. We are grateful to members of the Mellone lab for discussion and suggestions, especially Jason Palladino for early project design and members of the R. O’Neill lab for suggestions on genome analyses.

## References

Altemose N, Logsdon GA, Bzikadze AV, et al (2022) Complete genomic and epigenetic maps of human centromeres. Science 376:eabl4178. 10.1126/science.abl4178

Barnhart MC, Kuich PHJL, Stellfox ME, et al (2011) HJURP is a CENP-A chromatin assembly factor sufficient to form a functional de novo kinetochore. J Cell Biol 194:229–243. 10.1083/jcb.201012017

Baum M, Ngan VK, Clarke L (1994) The centromeric K-type repeat and the central core are together sufficient to establish a functional Schizosaccharomyces pombe centromere. Mol Biol Cell 5:747–761. 10.1091/mbc.5.7.747

Berger C, Renner S, Lüer K, Technau GM (2007) The commonly used marker ELAV is transiently expressed in neuroblasts and glial cells in the Drosophila embryonic CNS. Dev Dyn 236:3562–3568. 10.1002/dvdy.21372

Blower MD, Karpen GH (2001) The role of Drosophila CID in kinetochore formation, cell-cycle progression and heterochromatin interactions. Nat Cell Biol 3:730–739. 10.1038/35087045

Blower MD, Sullivan BA, Karpen GH (2002) Conserved Organization of Centromeric Chromatin in Flies and Humans. Dev Cell 2:319–330. 10.1016/s1534-5807(02)00135-1

Buenrostro JD, Wu B, Chang HY, Greenleaf WJ (2015) ATAC-seq: A Method for Assaying Chromatin Accessibility Genome-Wide. Curr Protoc Mol Biol 109:21.29.1-21.29.9. 10.1002/0471142727.mb2129s109

Chabot BJ, Sun R, Amjad A, et al (2024) Transcription of a centromere-enriched retroelement and local retention of its RNA are significant features of the CENP-A chromatin landscape. Genome Biol 25:295. 10.1186/s13059-024-03433-1

Chang C-H, Chavan A, Palladino J, et al (2019) Islands of retroelements are major components of Drosophila centromeres. Plos Biol 17:e3000241. 10.1371/journal.pbio.3000241

Chen C-C, Dechassa ML, Bettini E, et al (2014) CAL1 is the Drosophila CENP-A assembly factor. J Cell Biol 204:313–329. 10.1083/jcb.201305036

Deshmukh S, Ponnaluri VC, Dai N, et al (2018) Levels of DNA cytosine methylation in the Drosophila genome. PeerJ 6:e5119. 10.7717/peerj.5119

Dumont M, Fachinetti D (2017) Centromeres and Kinetochores, Discovering the Molecular Mechanisms Underlying Chromosome Inheritance. Prog Mol Subcell Biol 56:305–336. 10.1007/978-3-319-58592-5_13

Dunleavy EM, Roche D, Tagami H, et al (2009) HJURP Is a Cell-Cycle-Dependent Maintenance and Deposition Factor of CENP-A at Centromeres. Cell 137:485–497. 10.1016/j.cell.2009.02.040

Erhardt S, Mellone BG, Betts CM, et al (2008) Genome-wide analysis reveals a cell cycle–dependent mechanism controlling centromere propagation. J Cell Biol 183:805–818. 10.1083/jcb.200806038

Foltz DR, Jansen LET, Bailey AO, et al (2009) Centromere-Specific Assembly of CENP-A Nucleosomes Is Mediated by HJURP. Cell 137:472–484. 10.1016/j.cell.2009.02.039

Gershman A, Sauria MEG, Guitart X, et al (2022) Epigenetic patterns in a complete human genome. Science 376:eabj5089–eabj5089. 10.1126/science.abj5089

Giet R, Glover DM (2001) Drosophila Aurora B Kinase Is Required for Histone H3 Phosphorylation and Condensin Recruitment during Chromosome Condensation and to Organize the Central Spindle during Cytokinesis. J Cell Biol 152:669–682. 10.1083/jcb.152.4.669

Gratz SJ, Cummings AM, Nguyen JN, et al (2013) Genome Engineering of Drosophila with the CRISPR RNA-Guided Cas9 Nuclease. Genetics 194:1029–1035. 10.1534/genetics.113.152710

Guilinger JP, Thompson DB, Liu DR (2014) Fusion of catalytically inactive Cas9 to FokI nuclease improves the specificity of genome modification. Nat Biotechnol 32:577–582. 10.1038/nbt.2909

Hartley GA, Okhovat M, Hoyt SJ, et al (2025) Centromeric transposable elements and epigenetic status drive karyotypic variation in the eastern hoolock gibbon. Cell Genom 5:100808. 10.1016/j.xgen.2025.100808

Heeger S, Leismann O, Schittenhelm R, et al (2005) Genetic interactions of separase regulatory subunits reveal the diverged Drosophila Cenp-C homolog. Genes Dev 19:2041–2053. 10.1101/gad.347805

Hendzel MJ, Wei Y, Mancini MA, et al (1997) Mitosis-specific phosphorylation of histone H3 initiates primarily within pericentromeric heterochromatin during G2 and spreads in an ordered fashion coincident with mitotic chromosome condensation. Chromosoma 106:348–360. 10.1007/s004120050256

Henikoff S, Ahmad K, Malik HS (2001) The Centromere Paradox: Stable Inheritance with Rapidly Evolving DNA. Science 293:1098–1102. 10.1126/science.1062939

Hoyt SJ, Storer JM, Hartley GA, et al (2022a) From telomere to telomere: The transcriptional and epigenetic state of human repeat elements. Science 376:eabk3112. 10.1126/science.abk3112

Hoyt SJ, Storer JM, Hartley GA, et al (2022b) From telomere to telomere: The transcriptional and epigenetic state of human repeat elements. Science 376:eabk3112. 10.1126/science.abk3112

Jiang Y, Reichert H (2012) Programmed cell death in type II neuroblast lineages is required for central complex development in the Drosophila brain. Neural Dev 7:3–3. 10.1186/1749-8104-7-3

Kaya-Okur HS, Wu SJ, Codomo CA, et al (2019) CUT&Tag for efficient epigenomic profiling of small samples and single cells. Nat Commun 10:1930. 10.1038/s41467-019-09982-5

Logsdon GA, Ebert P, Audano PA, et al (2024) Complex genetic variation in nearly complete human genomes. bioRxiv 2024.09.24.614721. 10.1101/2024.09.24.614721

Madigan JP, Chotkowski HL, Glaser RL (2002) DNA double-strand break-induced phosphorylation of Drosophila histone variant H2Av helps prevent radiation-induced apoptosis. Nucleic Acids Res 30:3698–3705. 10.1093/nar/gkf496

Makova KD, Pickett BD, Harris RS, et al (2024) The complete sequence and comparative analysis of ape sex chromosomes. Nature 630:401–411. 10.1038/s41586-024-07473-2

Marshall OJ, Chueh AC, Wong LH, Choo KHA (2008) Neocentromeres: New Insights into Centromere Structure, Disease Development, and Karyotype Evolution. Am J Hum Genet 82:261–282. 10.1016/j.ajhg.2007.11.009

McKinley KL, Cheeseman IM (2016) The molecular basis for centromere identity and function. Nat Rev Mol Cell Bio 17:16–29. 10.1038/nrm.2015.5

Mehrotra S, McKim KS (2006) Temporal Analysis of Meiotic DNA Double-Strand Break Formation and Repair in Drosophila Females. PLoS Genet 2:e200. 10.1371/journal.pgen.0020200

Mendiburo MJ, Padeken J, Fülöp S, et al (2011) Drosophila CENH3 Is Sufficient for Centromere Formation. Science 334:686–690. 10.1126/science.1206880

Murillo-Pineda M, Valente LP, Dumont M, et al (2021) Induction of spontaneous human neocentromere formation and long-term maturation. J Cell Biology 220:e202007210. 10.1083/jcb.202007210

Musacchio A, Salmon ED (2007) The spindle-assembly checkpoint in space and time. Nat Rev Mol Cell Biol 8:379–393. 10.1038/nrm2163

Ohzeki J, Nakano M, Okada T, Masumoto H (2002) CENP-B box is required for de novo centromere chromatin assembly on human alphoid DNA. J Cell Biology 159:765–775. 10.1083/jcb.200207112

Palladino J, Chavan A, Sposato A, et al (2020) Targeted De Novo Centromere Formation in Drosophila Reveals Plasticity and Maintenance Potential of CENP-A Chromatin. Dev Cell 52:379–394.e7. 10.1016/j.devcel.2020.01.005

Palmer DK, O’Day K, Trong HL, et al (1991) Purification of the centromere-specific protein CENP-A and demonstration that it is a distinctive histone. Proc National Acad Sci 88:3734–3738. 10.1073/pnas.88.9.3734

Peng JC, Karpen GH (2009) Heterochromatic Genome Stability Requires Regulators of Histone H3 K9 Methylation. PLoS Genet 5:e1000435. 10.1371/journal.pgen.1000435

Port F, Bullock SL (2016) Augmenting CRISPR applications in Drosophila with tRNA-flanked sgRNAs. Nat Methods 13:852–854. 10.1038/nmeth.3972

Salinas-Luypaert C, Dubocanin D, Lee RJ, et al (2025) DNA methylation influences human centromere positioning and function. Nat Genet 57:2509–2521. 10.1038/s41588-025-02324-w

Scelfo A, Angrisani A, Grillo M, et al (2024) Specialized replication mechanisms maintain genome stability at human centromeres. Mol Cell 84:1003–1020.e10. 10.1016/j.molcel.2024.01.018

Scott KC, Merrett SL, Willard HF (2006) A Heterochromatin Barrier Partitions the Fission Yeast Centromere into Discrete Chromatin Domains. Curr Biol 16:119–129. 10.1016/j.cub.2005.11.065

Shang W-H, Hori T, Martins NMC, et al (2013) Chromosome Engineering Allows the Efficient Isolation of Vertebrate Neocentromeres. Dev Cell 24:635–648. 10.1016/j.devcel.2013.02.009

Shukla M, Tong P, White SA, et al (2018) Centromere DNA Destabilizes H3 Nucleosomes to Promote CENP-A Deposition during the Cell Cycle. Curr Biol 28:3924–3936.e4. 10.1016/j.cub.2018.10.049

Sullivan BA (2010) Fluorescence in situ Hybridization (FISH), Protocols and Applications. Methods Mol Biology 659:99–115. 10.1007/978-1-60761-789-1_7

Talbert PB, Masuelli R, Tyagi AP, et al (2002) Centromeric Localization and Adaptive Evolution of an Arabidopsis Histone H3 Variant. Plant Cell 14:1053–1066. 10.1105/tpc.010425

Ventura M, Antonacci F, Cardone MF, et al (2007) Evolutionary Formation of New Centromeres in Macaque. Science 316:243–246. 10.1126/science.1140615

Yoo D, Rhie A, Hebbar P, et al (2025) Complete sequencing of ape genomes. Nature 641:401–418. 10.1038/s41586-025-08816-3

